# Selective signatures and high genome-wide diversity in traditional Brazilian manioc (*Manihot esculenta* Crantz) varieties

**DOI:** 10.1101/2021.09.06.459170

**Authors:** Alessandro Alves-Pereira, Maria I. Zucchi, Charles R. Clement, João P. G. Viana, José B. Pinheiro, Elizabeth A. Veasey, Anete P. Souza

**Affiliations:** Departamento de Biologia Vegetal, Instituto de Biologia, Universidade Estadual de Campinas (UNICAMP). Av. Cândido Rondon, 400, Cidade Universitária, 13083-875, CP: 6010, Campinas, SP, Brazil; Centro de Biologia Molecular e Engenharia Genética, Universidade Estadual de Campinas (UNICAMP). Av. Cândido Rondon, 400, Cidade Universitária, 13083-875, CP: 6010, Campinas, SP, Brazil; Agência Paulista de Tecnologia dos Agronegócios (APTA), Pólo Centro-Sul. Rodovia SP 127, km 30, 13400-970, Piracicaba, SP, Brazil; Instituto Nacional de Pesquisas da Amazônia (INPA). Av. André Araújo, 2936, Petrópolis, 69067-375, Manaus, AM, Brazil; Department of Crop Sciences, University of Illinois at Urbana-Champaign (UIUC). AW-101 Turner Hall, 1102 South Goodwin Avenue, 61801-4798, Urbana, IL, USA; Departamento de Genética, Escola Superior de Agricultura “Luiz de Queiróz”, Universidade de São Paulo (ESALQ/USP). Av. Pádua Dias, 11, 13400-970, Piracicaba, SP, Brazil

**Author notes:** Corresponding author: Anete P. Souza.

**Keywords:** cassava, genetic resources, genotyping-by-sequencing, population genomics, SNP

## Abstract

Knowledge about crops’ genetic diversity is essential to promote effective use and conservation of their genetic resources, because genetic diversity enables farmers to adapt their crops to specific needs and is the raw material for breeding efforts. Currently, manioc (*Manihot esculenta* ssp. *esculenta*) is one of the most important food crops in the world and has the potential to help achieve food security in the context of on-going climate changes. In this study we assessed the patterns of genome-wide diversity of traditional Brazilian manioc varieties conserved in the gene bank of the Luiz de Queiroz College of Agriculture, University of São Paulo. We used single nucleotide polymorphisms to evaluate the organization of genetic diversity and to identify selective signatures contrasting varieties from different biomes with samples of manioc’s wild relative *M. esculenta* ssp. *flabellifolia*. We identified signatures of selection putatively associated with resistance genes, plant development and response to abiotic stresses. This presumed adaptive variation might have been important for the initial domestication and for the crop’s diversification in response to cultivation in different environments. The neutral variation revealed high levels of genetic diversity within groups of varieties from different biomes and low to moderate genetic divergence among biomes. These results reflect the complexity of manioc’s biology and its evolutionary dynamics under traditional cultivation. Our results exemplify how the smallholder practices contribute to the conservation of manioc’s genetic resources, maintaining variation of potential adaptive significance and high levels of neutral genetic diversity.

## Introduction

Food security – the regular access to enough high-quality food with sufficient protein and energy – is one of the major goals of the United Nations’ 2030 Agenda for Sustainable Development to promote a fairer world (United Nations 2015). It is, however, an enormous challenge due to the accelerated increase in the world’s human population and on-going global climate changes (Godfray et al. 2010; FAO 2021). Better strategies to conserve and use crop diversity are effective ways to address this issue, and, although often regarded as descriptive, the study of genetic diversity is fundamental to this end (Gepts 2006; FAO 2010).

Genetic diversity is used by farmers to adapt their crops to current and future climate changes and is also the raw material of formal breeding (Gepts 2006; McCouch et al. 2013). The maintenance of agrobiodiversity provides humans with a wealth of intraspecific crop diversity selected in different cultural and geographical contexts (Esquinas-Alcázar 2005; Castañeda-Álvarez et al. 2016). This agrobiodiversity is associated with valuable traditional knowledge and practice systems, which play a key role to conserve biological and cultural diversity (Fernández-Llamazares et al. 2021). Additionally, the management of landraces with high genetic diversity by farming communities has collaborated to keep reasonable levels of food production in many areas of the developing world (Esquinas-Alcázar 2005). However, factors such as globalization, degradation of natural landscapes, changes in agricultural production systems, and landrace displacement by modern cultivars threaten the conservation of plant genetic resources (Esquinas-Alcázar 2005; Gepts 2006).

Manioc (*Manihot esculenta* Crantz), also known as cassava, is currently one of the major food crops, ranking eighth in estimated global production (FAOSTAT 2019), and its roots are the main source of energy for more than 800 million people (Lebot 2009). The crop has an immense diversity of varieties which are cultivated around the Tropics, mainly by low-income smallholder farmers (Howeler et al. 2013; McKey and Delêtre 2017), and it is considered one of the most promising crops to promote food security in developing countries (Howeler et al. 2013). This is because manioc is well-adapted to marginal areas with poor soils and can be produced efficiently even on small scales, with low inputs and without mechanization (Howeler et al. 2013).

Morphological and genetic evidence support the hypothesis that manioc was domesticated from *M. esculenta* ssp. *flabellifollia* in southwestern Amazonia (Allem 1994; Olsen and Schaal 1999, 2001; Olsen 2004; Léotard et al. 2009; Mühlen et al. 2019), although there is still some controversy (Lebot 2009). Domestication may have started as early as 10,000 years before present (ybp) (Olsen and Schaal 1999), in what is now the Brazilian state of Rondônia and adjacent regions. Populations of ssp. *flabellifolia* currently occur in the Amazonia-Cerrado ecotones of southern Amazonia, as well as on the Guiana shield (Allem 1994). The wild progenitor grows as highly branched shrubs in open vegetation or as climbing vines amid denser vegetation, while cultivated manioc grows as little-branched shrubs (Ménard et al. 2013). After its initial domestication, manioc was probably available in most parts of the Neotropics by 6,500 ybp (Isendahl 2011; Brown et al. 2013). Manioc started to be globally spread after the European conquest of South America in the 16^th^ century, which introduced the crop in Africa, tropical Asia and Oceania (McKey and Delêtre 2017).

Cultivated manioc has two major groups of landraces that differ in their contents of cyanogenic compounds (in the whole plant, but especially in the edible roots). Sweet manioc has lower cyanogenic potential (< 100 ppm fresh weight) than bitter manioc (> 100 ppm fresh weight), but the variation in the content of cyanogenic compounds is continuous across these two groups (McKey et al. 2012). Sweet manioc can be safely consumed after simple processing (e.g., peeling and cooking), while bitter manioc needs more elaborate detoxification (e.g., peeling, soaking, grating, and roasting) (McKey and Beckerman 1993). Although sweet and bitter manioc cannot be separated by morphological traits, they are genetically divergent and farmers are able to recognize them (McKey and Delêtre 2017).

Manioc is a clonal crop, but its evolutionary dynamics is much more complex than the establishment of varieties consisting of unique clones. Under traditional cultivation, farmers propagate manioc varieties by stem-cuttings, but the plants can produce flowers (Lebot 2009). Manioc is allogamous, and crossings may occur between plants from different varieties producing fruits that disperse sexual seeds in the swiddens (McKey and Delêtre 2017). The seeds have elaiosomes that attract ants, which further disperse and bury them (Elias and McKey 2000). The sexual seeds in the soil seed banks may sprout when the swidden is cleared, or the vegetation of an abandoned field is burnt to start a new cycle of cultivation (Duputié et al. 2009). Farmers may consciously or inadvertently let sexual seedlings grow in the swiddens until harvesting time, when they may decide to use stem-cuttings of sexual plants for clonal propagation (Elias et al. 2001; Peroni et al. 2007). Farmers may either incorporate these stem-cuttings into an existing variety or start a new variety, increasing the crop’s genetic diversity (Martins 2007). This management of high genetic diversity extends beyond family units into extensive exchange networks of cultivated varieties (Coomes 2010; Dyer et al. 2011). These networks vary according to the cultural context, but are a common feature of traditional cultivation in Amazonia and throughout the world (Salick et al. 1997; Sambatti et al. 2001; Heckler and Zent 2008; Delêtre et al. 2011). By exchanging varieties that were selected and managed in the same or different regions, traditional farmers maintain high levels of genetic diversity in the crop at both local and wider geographical scales (Sardos et al. 2008; Delêtre et al. 2011).

The evolutionary dynamics outlined above is typical of manioc cultivation in Amazonia, but the crop’s dispersals around the world were not always accompanied by cultural appropriation, i.e., not all the original aspects of its cultivation can be observed outside Amazonia (McKey and Delêtre 2017). In the Neotropics, bitter landraces predominate where manioc is the major staple crop cultivated in swiddens far from household units, while sweet landraces predominate where manioc is part of multi-crop systems and is often cultivated in backyards (McKey and Delêtre 2017). These same patterns may be observed in most parts of Africa, but are much more variable in Asia and Oceania (Sardos et al. 2008; Ellen and Soselisa 2012). Farmer’s interest in experimenting with sexual seedlings also seems to vary greatly across locations outside the Neotropics (McKey and Delêtre 2017). Variable degrees of cultural appropriation may also influence the farmers’ ability to properly detoxify bitter manioc before safe consumption, which contributes to epidemics of *Konzo*, a chronic paralytic disease, in some African regions (McKey and Delêtre 2017). These differences in the crop’s management influence how manioc evolves under distinct geographical and cultural contexts and have a significant impact in the crop’s potential to contribute to food security (Burns et al. 2010).

Genetic studies have greatly contributed to our understanding about the evolution of the crop (Elias et al. 2001; Sambatti et al. 2001; Pujol et al. 2005a) and to support breeding (Mba et al. 2001; Oliveira et al. 2012; Ferguson et al. 2019). More recently, approaches assessing genome-wide diversity of domesticated varieties and wild relatives (Bredeson et al. 2016; Kuon et al. 2019; Wolfe et al. 2019) advanced rapidly after the release of a genome for manioc (Prochnik et al. 2012). Indeed, the characterization of genome-wide diversity collaborated tremendously for the valorization and utilization of wild species and cultivated varieties conserved in gene banks throughout the world (Rabbi et al. 2015; Albuquerque et al. 2018; Ogbonna et al. 2021b). Given the agronomic importance of manioc, these studies were mostly developed to support breeding. However, the genomic approaches also offer opportunities to further investigate evolutionary aspects related to the crop’s domestication and diversification (Morrell et al. 2012; Allendorf 2017).

In this context, we assessed selective signatures and the genome-wide diversity of manioc varieties from different Brazilian biomes, based on single nucleotide polymorphisms (SNP). The varieties are conserved at the Luiz de Queiroz College of Agriculture gene bank, Brazil, and are compared with wild samples collected in the center of manioc domestication. We performed genome scans to identify SNPs with putative signatures of selection and discuss their potential relevance to the domestication and diversification processes associated with cultivation in different environmental contexts. We aimed to generate novel information about the evolution of manioc in its country of origin and to contribute to better management of the genetic resources of this globally important crop.

## Material and Methods

### Plant material and DNA isolation

We sampled apical leaves of the 78 manioc varieties (1 plant/accession) from the gene bank at the Luiz de Queiroz College of Agriculture, University of São Paulo, Piracicaba, São Paulo, Brazil (22°42’26,8” S; 47°38’17,8” W). These varieties were originally collected in smallholder farmer communities in six Brazilian states, located in four different biomes: Amazonia (16), Cerrado (30), Atlantic Forest (27) and Pantanal (5) (Figure 1, Table S1). We also evaluated other samples collected in a previous study (Alves-Pereira et al. 2020): eight *M. esculenta* ssp. *flabellifolia* samples collected in the center of manioc domestication in Rondônia state, and six Amazonian varieties (three bitter and three sweet). We used these samples as references for the crop’s closest wild relative, and for the major groups of cultivated manioc, respectively. The collections of these samples were registered in CGEN (numbers A7994B4 and AEA71DE), according to Brazilian Law 13123 (20 May 2015).

**Figure 1.**
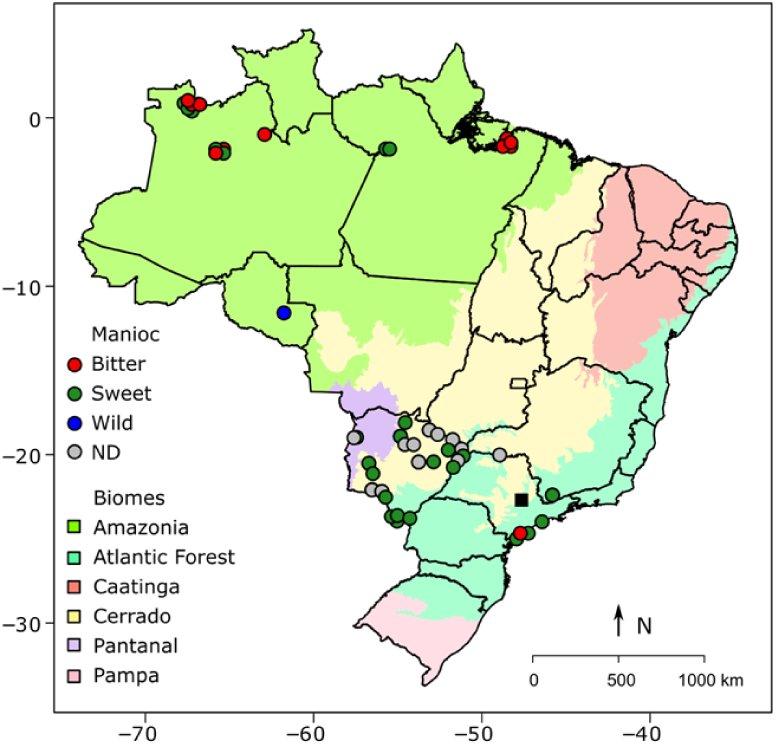
Map of Brazil showing the geographical locations of the municipalities in which manioc (*Manihot esculenta*) samples were originally collected (some points were slightly moved for easier visualization). The black square indicates the location of the gene bank at the Luiz de Queiroz College of Agriculture, Piracicaba, São Paulo, Brazil. ND = no data on toxicity.

The leaves were dehydrated with silica gel in paper bags and stored at -20°C. We obtained total genomic DNA from 50 mg of leaf samples following the protocol described by Doyle and Doyle (1987) and inspected DNA quality with electrophoresis in agarose 1 % (w/v) gels stained with ethidium bromide. We estimated DNA concentrations with dsDNA BR Assay quantification kit for Qubit3 fluorometer (Invitrogen), and normalized DNA concentrations to 25 ng·uL^-1^.

### Genomic libraries and SNP identification

We prepared two double-digest genotyping-by-sequencing (ddGBS) libraries as described by Poland et al. (2012). Briefly, 175 ng of genomic DNA were digested with *PstI* and *MseI* for one library, and *NsiI* and *MspI* for the other (all enzymes from New England Biosciences). The restriction fragments were ligated to adaptors complimentary to each of the restriction sites (including 96-plex *PstI* or *NsiI* adaptor sets with unique 4-9 bp barcode sequences). Ligation products of each sample were pooled and enriched for fragments containing different adapters in both ends through PCR. The concentrations of the ddGBS libraries were estimated using the NEBNext Library Quant kit for Illumina (New England Biosciences) through real-time PCR, and the libraries’ profiles were inspected using the DNA 12000 Analysis kit for Bioanalyzer 2100 (Agilent). Each ddGBS library was sequenced twice in the NextSeq550 (Illumina) platform (single-end, 150 bp).

We inspected the quality of raw reads using FastQC (Andrews 2010), and due to a large amount of 3’-end adaptors we trimmed the reads to 100 and 80 bp for *PstI*-*MseI* and *NsiI*-*MspI* libraries, respectively. We performed read trimming and demultiplex according to specific barcodes using the module *process_radtags* of Stacks 1.42 (Catchen et al. 2013). We aligned the demultiplexed reads against the manioc genome *Manihot esculenta v6* (NCBI PRJNA234389) (Prochnik et al. 2012) using Bowtie 2.2.1 (Langmead and Salzberg 2012) with the “end-to-end” and “sensitive” configurations. We identified SNPs separately for each library using SAMtools 0.1.19 (Li et al. 2009; Li 2011) and VCFtools 0.1.17 (Danecek et al. 2011), retaining only biallelic markers and only one SNP per read to avoid explicit linkage. Candidate SNPs had sequence depth ≥ 5X, minor allele frequency ≥ 0.05, mapping quality ≥ 13 and were present in at least 90 % of the samples. The SNPs identified in each library were merged using MergeVcfs from Picard (http://broadinstitute.github.io/picard). Then, we used VCFtools 0.1.17 (Danecek et al. 2011) and SAMtools 0.1.19 (Li 2011) to filter out SNPs within 100 bp and within windows of 1,000 SNP sites with linkage disequilibrium (LD) r^2^ ≥ 0.8, and to retain the SNPs observed in *Manihot esculenta v6* chromosomes.

### Genome scans

We used five different methods (*pcadapt, F*_*ST*_, *FLK, hapFLK*, and *XP-EHH*) with distinct underlying models to detect putative selective signatures (outlier SNPs). These analyses were performed considering two scenarios: i) the groups of wild versus cultivated manioc, and ii) the groups of varieties per biomes (without wild samples). With this design we aimed to identify loci with putative selective signatures related to either the domestication of manioc or the diversification of the crop in different cultivation environments.

Pcadapt is based on a principal component analysis (PCA) without assuming any explicit genetic model (Luu et al. 2017). The detected outlier SNPs are associated with the first *K* principal components with greater contribution to the observed genetic structure. We used pcadapt 4.03 (Luu et al. 2017) for R (R Core Team 2018) to test 1 to 20 *K* principal components. We detected outliers based on *K* = 2 for wild versus cultivated manioc, and *K* = 5 for the biomes (Figure S1). The estimation of genetic differentiation among populations, measured by *F*_*ST*_ or related statistics, is a classic method for detecting selective signatures that reflect a broad range of scenarios, such as selection of standing variation or incomplete sweeps (Fariello et al. 2013). We estimated Weir and Cockerham’s *F*_*ST*_ (1984) among the groups of samples for each locus using diveRsity (Keenan et al. 2013) for R (R Core Team 2018). *FLK* is similar to *F*_*ST*_, but it accounts for hierarchical structures using a kinship matrix to model the covariance of the populations’ allele frequencies (Bonhomme et al. 2010). We estimated *FLK* for each locus using hapFLK 1.4 (https://forge-dga.jouy.inra.fr/projects/hapflk) based on the calculation of Reynold’s (1983) distances from our data.

The three methods above are based on individual markers, and we also applied two tests (*hapFLK* and *XP-EHH*) considering “long-range haplotypes” that account for variation in recombination rates by comparing haplotypes to other alleles in adjacent loci (Sabeti et al. 2007). *HapFLK* was built upon *FLK* for the detection of positive selection, including incomplete sweeps, from multiple populations and is robust in the presence of bottlenecks and migration (Fariello et al. 2013). Complimentarily, the cross-population extended haplotype homozygosity (*XP-EHH*) detects loci that have swept to near fixation (hard sweeps) within a specific population (Sabeti et al. 2007). For that reason, this analysis was performed only considering the groups of cultivated and wild manioc. For these two analyses, we used fastPHASE 1.4 (Scheet and Stephens 2006) to estimate missing data and reconstruct haplotypes. We used a perl script (https://github.com/lstevison/vcf-conversion-tools) to convert VCF files to fastPHASE input format and then performed 20 runs of the expectation-maximization (EM) algorithm with default configurations. We estimated *hapFLK* simultaneously with *FLK* considering the same Reynold’s distance matrix and using 15 EM runs. We estimated *XP-EHH* using the functions *scan_hh(), ies2xpehh()* and *calc_candidate_regions()* in rehh 2.0 (Gautier et al. 2017) for R (R Core Team 2018). We set the functions to use unpolarized data, and to consider the XP-EHH estimates in the direction of wild to cultivated manioc (positive selection of alleles in the cultivated samples).

We considered as outlier SNPs the loci with *q*-values ≤ 0.10 in *pcadapt*, loci at the top and bottom 2.5 % of the *F*_*ST*_ estimates (reflecting positive and balancing selection, respectively), and the loci at the top 5 % of the *FLK, hapFLK* and *XP-EHH* estimates. Then, we considered as loci putatively under selection those SNPs that were detected in at least two of the five (four) methods used for the groups of wild versus cultivated manioc (groups of varieties per biome).

We evaluated the predicted effects of outlier SNPs using SnpEff (Cingolani et al. 2012). We recovered the Gene Ontology (GO) annotations, which summarize the information about biological processes, molecular functions, and cellular components (Blake 2013) of the gene sequences with outlier SNPs extracted from *Manihot esculenta v6* with BEDTools 2.30.0 (Quinlan and Hall 2010). Genes with outlier SNPs were tested for enrichment of gene function descriptions using topGO 2.44.0 (Alexa and Rahnenführer 2021) for R (R Core Team 2018) based on default configurations and a threshold of p < 0.01 for Fisher’s exact tests. We compared the amino acid sequences of the predicted manioc genes with i) the predicted protein sequences of 2,609 manioc R-genes from the Plant Resistance Gene database (PRGdb 3.0) (Osuna-Cruz et al. 2018); and ii) the Swiss-Prot database from UniProt (www.uniprot.org) using blastp from blast 2.7.1 (Camacho et al. 2009). Then, we further assessed putative functional annotations described in UniProt for the blastp hits with the arbitrary threshold of identity ≥ 90 % (PRGdb) or ≥ 60 % (Swiss-Prot).

### Population genetics analyses

We performed the analyses of genetic diversity and structure considering only the putatively neutral SNPs (those that were not detected as outliers by at least two of the tests described above). We estimated the genetic diversity [total number of alleles (*A*), percentage of polymorphic loci (*%P*), number of private alleles (*PA*), observed (*H*_*O*_) and expected (*H*_*E*_) heterozygosities] and the inbreeding coefficients (*f*) using diveRsity (Keenan et al. 2013) and PopGenKit (Paquette 2012) for R (R Core Team 2018). *H*_*O*_, *H*_*E*_, and *f* confidence intervals were obtained with 1,000 bootstraps.

We evaluated the genetic variation within and among groups of varieties with analyses of molecular variance (AMOVA), estimated pairwise genetic divergence (F_ST_) among groups, and obtained their associated significance based on 20,000 permutations using Arlequin 3.5 (Excoffier and Lischer 2010). We investigated the patterns of genetic structure using sparse non-negative matrix factorization (sNMF) (Frichot et al. 2014) and discriminant analysis of principal components (DAPC) (Jombart et al. 2010). sNMF assumes that the genetic data originate from the admixture of *K* unknown parental populations and estimates ancestry coefficients from multilocus genotypes (Frichot et al. 2014). This analysis is similar to other model-based approaches, such as Structure (Pritchard et al. 2000), with the advantage of being robust to departures from traditional population genetic model assumptions (Frichot et al. 2014). We tested from 1 to 10 *K* ancestral populations with 200,000 iterations, ten repetitions for each *K* value, and used the cross-entropy criterion to visualize the results of the simulations. We performed this analysis using LEA (Frichot and François 2015) for R (R Core Team 2018). Complimentarily, DAPC summarizes information from large datasets (like genotypes from thousands of SNPs) to assign individuals to clusters without a pre-defined genetic model (Jombart et al. 2010). DAPC is useful for the assessment of genetic structure based on SNP datasets because it maximizes the variation among groups while minimizing the correlations between the original variables (such as LD) (Jombart et al. 2010). We performed DAPCs in adegenet (Jombart and Ahmed 2011) for R (R Core Team 2018) based on the groups of wild and cultivated manioc, and the groups of varieties per biomes (without wild samples).

## Results

### SNP genotyping

Sequencing of the two ddGBS libraries resulted in 198,017,706 (*NsiI*-*MspI*) and 240,176,492 (*PstI*-*MseI*) raw reads. After demultiplexing and quality filtering, 153,858,282 reads for *NsiI*-*MspI* (mean of 1,672,372.6 reads per sample ± 406,184.6 SD) and 118,476,082 reads for *PstI*-*MseI* (mean of 1,287,783.5 reads per sample ± 540,792.9 SD) were used for SNP identification. The *NsiI*-*MspI* library resulted in 4,790 SNPs (34.8 mean sequence depth per locus ± 21.3 SD, 2.2 % of missing data) while the *PstI*-*MseI* library resulted in 10,121 SNPs (29.5 mean sequence depth per locus ± 16.8 SD, 4.8 % of missing data). After merging these data sets and pruning markers with high LD we obtained a final set of 11,782 high-quality SNPs (31.2 mean sequence depth per locus ± 18.5 SD, 4 % of missing data) (Figure S2, Table S2).

### Putative signatures of selection

The tests for selective signatures contrasting the wild and cultivated manioc identified a total of 2,301 outlier SNPs (*pcadapt*: 1,239; *F*_*ST*_: 588; *FLK*: 590; *hapFLK*: 590; *XP-EHH*: 364), of which 698 SNPs were identified by at least two different methods (Figures 2A, 2C). When contrasting the groups of varieties per biome, we identified a total of 1,673 outlier SNPs (*pcadapt*: 161; *F*_*ST*_: 550; *FLK*: 556; *hapFLK*: 590), of which 169 SNPs were identified by at least two different methods (Figures 2B, 2D). Only two outlier SNPs were common for both criteria, and we considered that 865 outlier SNP loci showed putative signatures either of the selection of cultivated manioc from the wild ancestor, or for diversification of manioc in different cultivation environments. A total of 5,174 effects were predicted for these outlier SNPs (Table S3), of which 569 were within introns and 534 within exons (269 synonymous mutations and 265 non-synonymous). These numbers are greater than the number of outlier SNPs because the effects were predicted for all the alternative transcripts of the genes.

**Figure 2.**
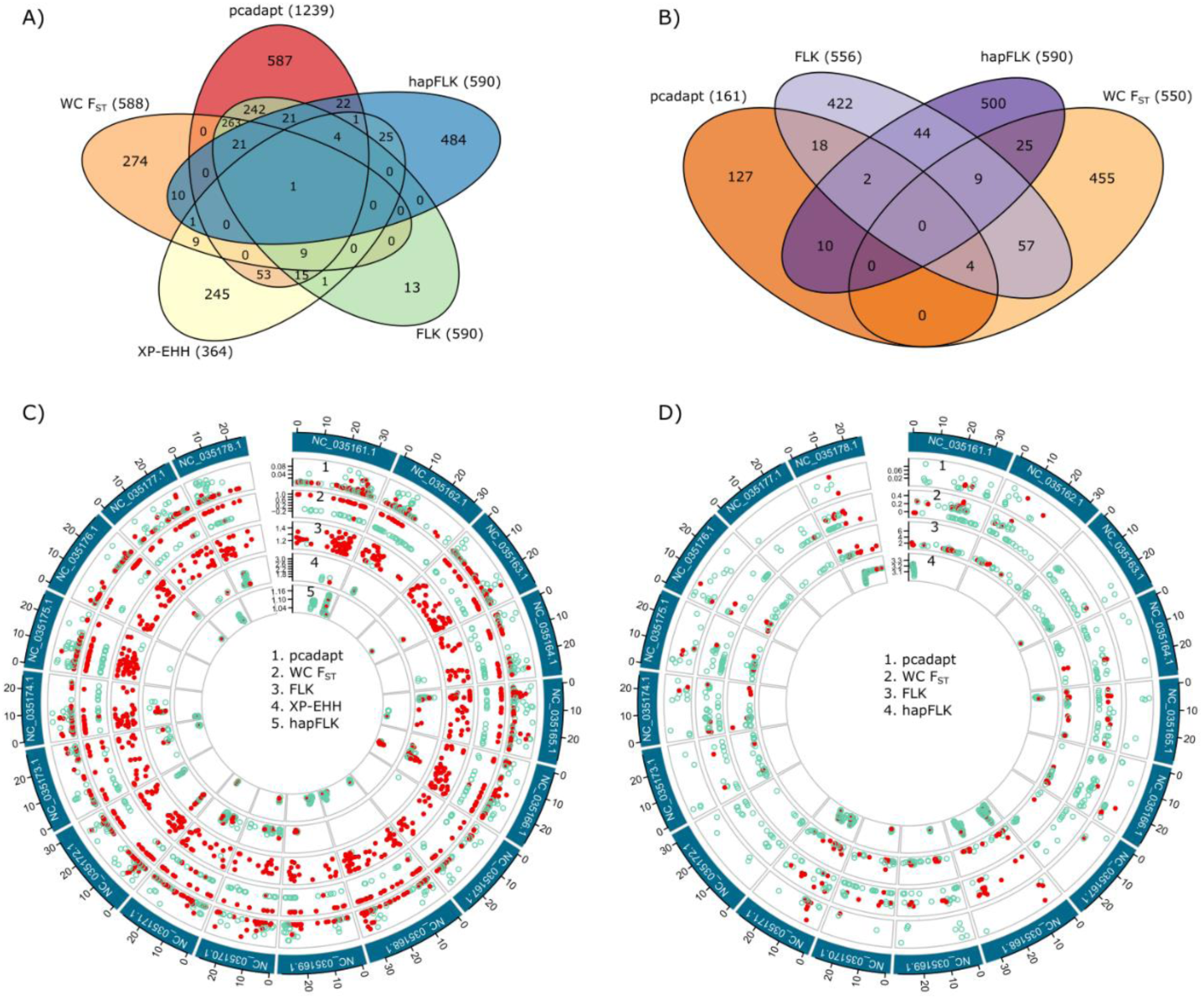
Summary of genome scans for signatures of selection considering different groups of manioc (*Manihot esculenta*) samples. Venn diagrams showing the number of outlier SNPs detected for each test (within parenthesis) and the overlap among them (numbers inside ellipses) for A) wild and cultivated manioc, and B) the groups of varieties per biome. The genomic context of outlier SNPs is illustrated in circular plots for C) the groups of wild and cultivated manioc, and D) the groups of varieties per biome. Each manioc chromosome is represented by a different box (10 Mb tick sizes), and their names are coded according to the manioc genome *Manihot esculenta v6* (NCBI PRJNA234389). The outlier SNPs are represented by dots for each test, which are shown in different layers. The outlier SNPs detected by at least two tests are highlighted in red.

Among the loci with putative selective signatures, 680 SNPs were in 663 different predicted manioc genes. These genes showed a total of 1,176 annotations distributed in 33 different GO classes (Figure S3). The most frequent GO annotations were associated with the molecular functions of binding (215 genes) and catalytic activity (188 genes), and the biological process of metabolism (181 genes). Consistent with this, the most frequent enriched GO annotations were binding to ATP (58 genes) and biding to metal ion (49 genes) (Table S4). A total of 69 manioc genes with outlier SNPs were similar to PRGdb 3.0 resistance genes (45 with identity > 90 %) belonging to six different classes (according to their protein domains), and most of them (38) had a kinase domain. A total of 576 manioc genes with outlier SNPs were similar to Swiss-Prot proteins (306 with identity > 60 %) related to many functions, including plant growth and development, organ sizes, root, flowering, and biotic and/or abiotic stresses. Many genes had similarities with proteins related to more general cellular processes, such as cell proliferation/elongation, transcription regulation, chloroplast activity, signaling, and ubiquitination. The variety of functions is exemplified by the descriptions for 21 of these proteins (Appendix 1), that we used to guide our discussion. Table S5 summarizes blastp results.

### Genome-wide diversity

The analyses below were performed considering the 10,917 putatively neutral SNPs that were not identified as outliers by more than one test of selection. We report the results based on all the 92 samples evaluated, but similar results were observed when considering only the varieties conserved in the gene bank (Tables S6, S7 and S8). Likely due to the larger sampling number, cultivated manioc had greater allelic diversity, more private alleles, and higher expected heterozygosity than wild manioc (Table 1). Within cultivated manioc, the sweet varieties had greater observed than expected heterozygosity (*H*_*O*_ = 0.322, *H*_*E*_ = 0.308), while the bitter varieties had a slight deficit of heterozygotes (*H*_*O*_ = 0.295, *H*_*E*_ = 0.309), although its small positive inbreeding coefficient (*f* = 0.047) was not significant. The varieties from different biomes had similar levels of genetic diversity (Table 1). Amazonia was the only biome with a slight deficit of heterozygotes (*f* = 0.062), but it had the greatest number of private alleles (*PA* = 153).

**Table 1.**
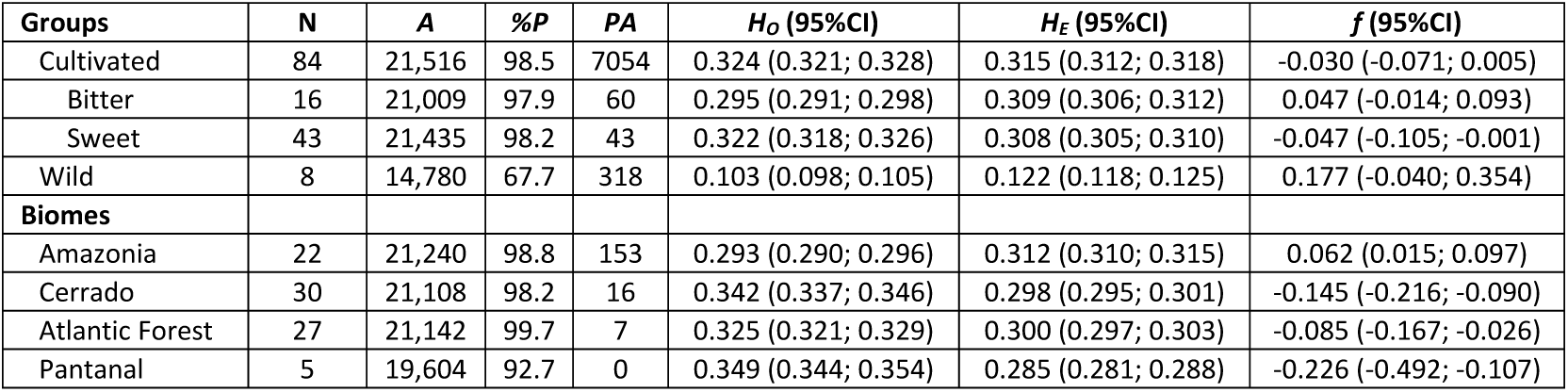
Estimates of genetic diversity and inbreeding based on 10,917 neutral SNPs for groups of manioc (*Manihot esculenta*) varieties. Number of samples (N), Total number of alleles (*A*), percentage of polymorphic loci (*%P*), number of private alleles (*PA*), observed (*H*_*O*_) and expected (*H*_*E*_) heterozygosities, inbreeding coefficients (*f*), and 95 % confidence intervals (95%CI).

According to the AMOVAs, greater proportions of the genetic variance were found within groups (Table 2). As expected, the greatest divergence among groups was observed between wild and cultivated manioc (ϕ_ST_ = 0.32, p < 0.001), followed by the divergence between the groups of biomes when compared to wild manioc (ϕ_CT_ = 0.31, p < 0.001). The divergence among different biomes was small, yet significant (ϕ_ST_ = 0.03, p < 0.001). Pair-wise estimates of *F*_*ST*_ between cultivated and wild manioc were high (Table 3). The bitter varieties were slightly more divergent from wild manioc (*F*_*ST*_ = 0.357) than the sweet varieties (*F*_*ST*_ = 0.344), and the divergence between bitter and sweet varieties was much lower (*F*_*ST*_ = 0.045), yet significant. All the biomes were highly divergent from wild manioc, with *F*_*ST*_ranging from 0.34 (Amazonia) to 0.43 (Pantanal). The divergence between biomes ranged from non-existent (Atlantic Forest vs. Pantanal = -0.015) to moderate (Amazonia vs. Cerrado = 0.065). Amazonia had the greater and significant estimates of divergence in relation to the other biomes, although *F*_*ST*_ was low to moderate (Table 3).

**Table 2.**
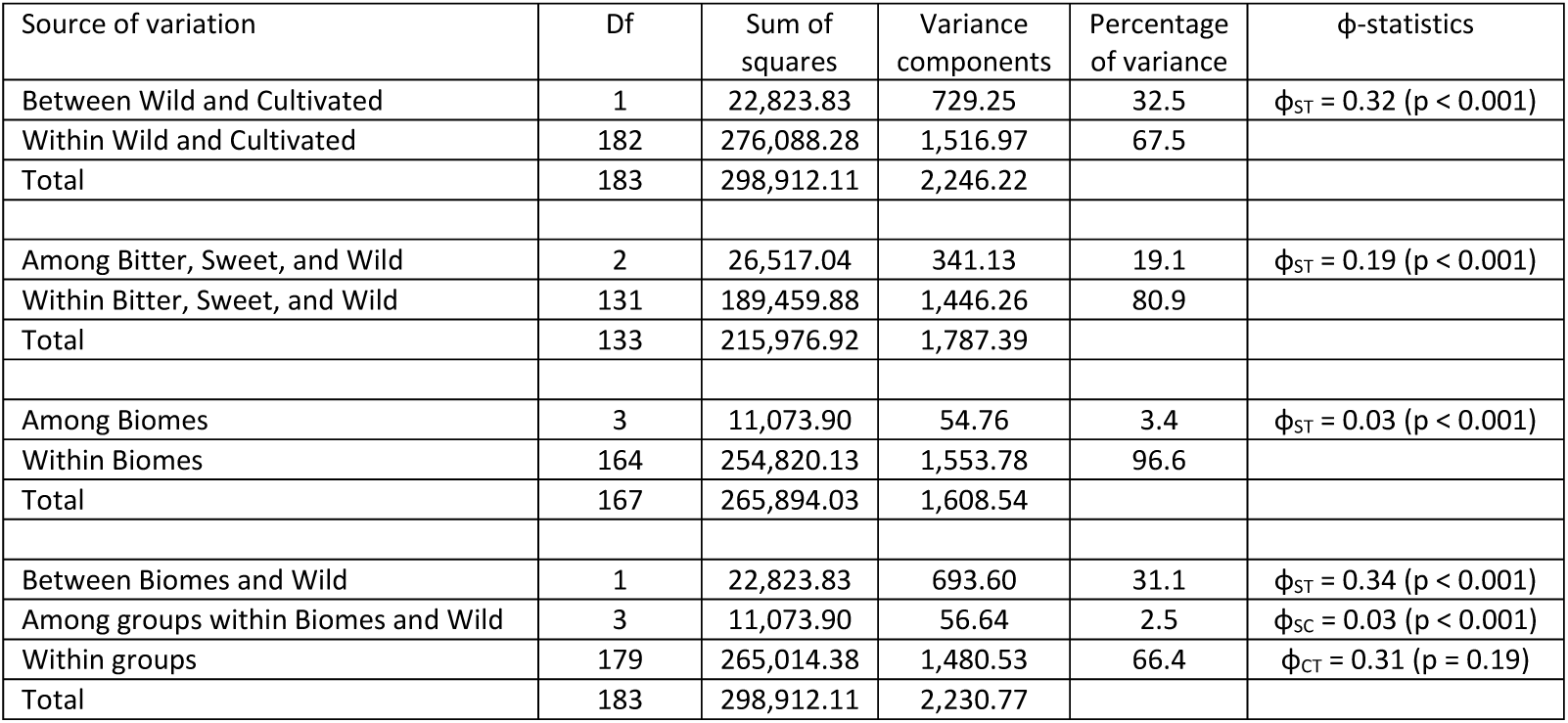
Analysis of molecular variance based on 10,917 neutral SNPs, showing the genetic variation within and among hierarchical groups of manioc (*Manihot esculenta*) varieties. Degrees of freedom (Df).

| Source of variation | Df | Sum of squares | Variance components | Percentage of variance | ϕ-statistics |
| --- | --- | --- | --- | --- | --- |
| Between Wild and Cultivated | 1 | 22,823.83 | 729.25 | 32.5 | $\phi_{ST} = 0.32$ ($p < 0.001$) |
| Within Wild and Cultivated | 182 | 276,088.28 | 1,516.97 | 67.5 |  |
| Total | 183 | 298,912.11 | 2,246.22 |  |  |
| Among Bitter, Sweet, and Wild | 2 | 26,517.04 | 341.13 | 19.1 | $\phi_{ST} = 0.19$ ($p < 0.001$) |
| Within Bitter, Sweet, and Wild | 131 | 189,459.88 | 1,446.26 | 80.9 |  |
| Total | 133 | 215,976.92 | 1,787.39 |  |  |
| Among Biomes | 3 | 11,073.90 | 54.76 | 3.4 | $\phi_{ST} = 0.03$ ($p < 0.001$) |
| Within Biomes | 164 | 254,820.13 | 1,553.78 | 96.6 |  |
| Total | 167 | 265,894.03 | 1,608.54 |  |  |
| Between Biomes and Wild | 1 | 22,823.83 | 693.60 | 31.1 | $\phi_{ST} = 0.34$ ($p < 0.001$) |
| Among groups within Biomes and Wild | 3 | 11,073.90 | 56.64 | 2.5 | $\phi_{SC} = 0.03$ ($p < 0.001$) |
| Within groups | 179 | 265,014.38 | 1,480.53 | 66.4 | $\phi_{CT} = 0.31$ ($p = 0.19$) |
| Total | 183 | 298,912.11 | 2,230.77 |  |  |

**Table 3.** Pairwise estimates of genetic divergence (Weir & Cockerham’s *F*_*ST*_ (1984)) among groups of manioc (*Manihot esculenta*) varieties, based on 10,917 neutral SNPs. Asterisks indicate significant estimates at p < 0.01.

| Groups | Bitter | Sweet |  |  |
| --- | --- | --- | --- | --- |
| Bitter |  |  |  |  |
| Sweet | 0.045* |  |  |  |
| Wild | 0.357* | 0.344* |  |  |
| Biomes | Amazonia | Cerrado | Atlantic Forest | Pantanal |
| Amazonia |  |  |  |  |
| Cerrado | 0.065* |  |  |  |
| Atlantic Forest | 0.045* | 0.017 |  |  |
| Pantanal | 0.044* | -0.025 | -0.015 |  |
| Wild | 0.342* | 0.365* | 0.362* | 0.433* |

The high divergence between wild and cultivated manioc, and the low divergence among biomes were also evident in sNMF and DAPC (Figure 3). There was no flatting point in the curve of cross-entropy estimates in sNMF, suggesting the absence of major genetic structure (Figure 3A). Therefore, we evaluated the correspondence of the ancestry coefficients for *K* = 2, 3 and 5 with the respective groups of cultivated vs. wild, bitter vs. sweet vs. wild, and the four biomes vs. wild (Figure 3B). The most evident genetic structure was observed between the wild and cultivated manioc, with high admixture between the groups of bitter and sweet varieties or among biomes. DAPC results were similar, showing high divergence between wild and cultivated manioc, and a great overlap of varieties from different biomes (Figures 3C and 3D). Besides the great genetic admixture, the DAPC plots also identify some highly divergent varieties from different biomes. Because DAPC maximizes between-group variations, Amazonia was somewhat more divergent in relation to the other biomes, just as suggested by the pairwise *F*_*ST*_ estimates.

**Figure 3.**
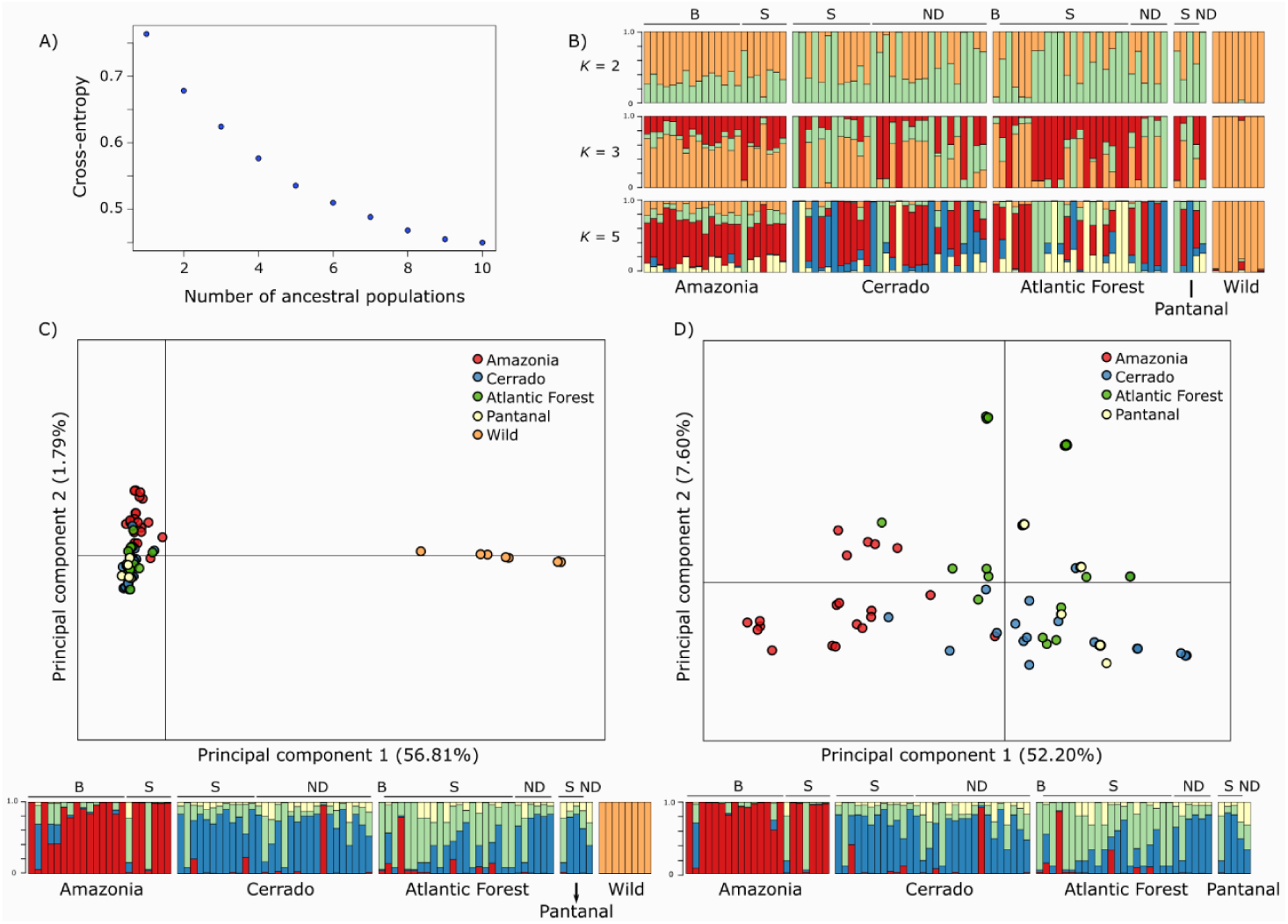
Genetic structure of 92 manioc (*Manihot esculenta*) varieties based on 10,917 neutral SNPs. A) Plot of cross-entropy estimates for different numbers of ancestral populations (*K*) in sparse non-negative matrix factorization (sNMF) showing no evident flatting point in the curve corresponding to the most-likely number of ancestral populations. B) Bar plots of sNMF ancestry coefficients for *K* = 2, 3, and 5. Discriminant analyses of principal components (DAPC) considering: C) the groups of wild manioc and the different biomes, and D) only the cultivated varieties grouped by biomes. The respective membership coefficients of each DAPC are shown as bar plots below scatter plots. Cultivated manioc is ordered in the bar plots according to the biomes and their reputed toxicity (B = bitter, S = sweet, ND = non-designated).

## Discussion

### Genome scans

There are many methods for the detection of selective signatures based on the significant deviation of outlier markers from the distribution of a given statistic measured under a specific model (Ahrens et al. 2018). However, deviance from model assumptions and covariance with sampling strategy, demography, underlying genetic structure, and other specific factors may lead to the detection of false positives (Lotterhos and Whitlock 2014, 2015; Hoban et al. 2016; Ahrens et al. 2018). A general approach to account for these limitations is to combine the results of different outlier tests (e.g., Pankin et al. 2018; Maccaferri et al. 2019). The variable number of outlier SNPs detected by the tests we performed reflect their different underlying models. We discuss below the possible biological significance of some outlier SNPs consistently identified by different tests.

The reduction of genetic diversity associated with domestication bottlenecks, or resulting from multiple founding effects during crops’ dispersals (Allaby et al. 2019; Brown 2019), might have affected plant defense and resistance mechanisms (Gaillard et al. 2018). The presence of outlier SNPs in putative resistance genes from different classes suggests that the gene bank conserves important genetic resources for the crop. For example, the outlier SNPs *Pst_1761* and *Pst_6563* were identified in genes similar to the disease resistance proteins RPS2 and RPM1, which are involved in the response to bacterial blight. In manioc, “cassava bacterial blight” is caused by *Xanthomonas campestris* pv. *manihotis* (or *X. axonopodis* pv. *manihotis*) that is found in cultivation areas throughout the world and is one of the most serious diseases affecting the crop (Hillocks and Wydra 2002; Lebot 2009). We also identified outlier SNPs (*Nsi_172* and *Pst_4209*) in genes similar to proteins involved in resistance to powdery mildew fungus, which in manioc is known as ash disease and is caused by *Oidium manihotis* Henn. (Hillocks and Wydra 2002). Although ash disease is widespread, it is not considered of great importance due to its superficial lesions (Hillocks and Wydra 2002). Agronomic trials would be important to confirm if some of the varieties conserved in the gene bank may be sources of resistance alleles for these diseases. Pest and disease resistance may be the principal weakness of manioc to adapt to climate change (Jarvis et al. 2012).

Most of the putative manioc resistance genes had other functions probably due to their catalytic kinase domains, which play many key roles in eukaryotes (Hanks 2003) including signaling and plant defense responses (Champion et al. 2004; Meng and Zhang 2013). Other outlier SNPs were in genes related to general cellular processes, such as ubiquitination (*Nsi_2393, Nsi_3361* and *Pst_4290*) and transcriptional regulation (*Nsi_3361, Pst_8996, Pst_9853* and *Pst_9959*). These processes are central in the expression and regulation of several other genes, and therefore are likely to be involved in adaptations, including domestication traits (Lenser and Theißen 2013; Gepts 2014).

Outlier SNPs in genes putatively involved with development may be genetic signatures of manioc domestication and selection for different traits of interest in response to distinct human preferences. Manioc was domesticated for its starchy roots, and we identified SNPs in genes putatively involved in root formation (*Nsi_2393, Nsi_3361, Pst_4290, Pst_8996* and *Pst_9959*), organ size (*Nsi_3361*) and shape (*Pst_1010*), and in starch metabolism (*Pst_5701*). These SNPs may be of agronomic importance because the current major objectives of breeding for food and industrial uses include increasing the root/stem ratio, improving starch quality, and increasing starch content in roots (Jennings and Iglesias 2002; Ceballos et al. 2007). We also found outlier SNPs in genes putatively involved in the development of shoots (*Nsi_3361 and Nsi_3540*) and branching (*Pst_6259*). Domestication often resulted in changes in shoot architecture that facilitate plant growth and harvesting (Meyer et al. 2012; Meyer and Purugganan 2013). Contrasting with the highly branched habit of ssp. *flabellifolia* (Ménard et al. 2013), farmers prefer sparsely branched manioc plants because they may provide thicker stem cuttings (Elias et al. 2007). Some outlier SNPs were in genes putatively involved in flower development (*Pst_7007, Pst_8971, Pst_9853* and *Pst_9959*), fertilization (*Nsi_172* and *Pst_4209*), and gametogenesis (*Nsi_3361*). These selective signatures may result from the relaxation of selective pressures on sexual fertility caused by domestication for vegetative propagation (Zohary 2004). Flowering is variable in manioc and about 60 % of the pollen produced may remain viable, although manioc cultivars commonly have male sterility and low seed/flower ratios (Lebot 2009). We also identified outlier SNPs putatively involved in cotyledon development (*Nsi_2289* and *Nsi_4509*); this was in a manioc gene similar to *Arabidopsis thaliana* STY46, a protein kinase involved in chloroplast biogenesis and differentiation in cotyledons (Lamberti et al. 2011). The seedlings of domesticated and wild manioc have contrasting functional differences (Pujol et al. 2005b). The cotyledons are hypogeal in the seedlings of ssp. *flabellifolia*, providing additional opportunities for plant regrowth after damage. In domesticated manioc, cotyledons are epigeal, foliaceus, and photosynthetically active, collaborating with rapid plant growth (Pujol et al. 2005b).

Growth resilience under distinct environments was essential for the spread and adaptation of crops to regions outside their domestication centers (Meyer and Purugganan 2013). The outlier SNP *Pst_29* was in a gene putatively involved in the metabolism of glycosinolates, secondary compounds that may act in plant defense to a wide range of enemies (Halkier and Gershenzon 2006). Ecological shifts to resource-richer cultivation environments may have relaxed the selection for chemical and physical defenses in some crops (McKey et al. 2012). For manioc, natural selection might have been important to maintain defense mechanisms because cultivation in anthropic landscapes made plants more apparent to pests and herbivores (McKey et al. 2012). This is especially important in the context of active human selection for low toxic sweet manioc and cultivation in herbivore and pathogen-rich tropical regions (McKey et al. 2012), such as Amazonia and other Brazilian biomes. We also found outlier SNPs (*Nsi_172, Pst_1010, Pst_3333, Pst_4209* and *Pst_5985*) possibly involved in cell proliferation and elongation. Genes controlling cell division are frequently associated with domestication and diversification processes because they often resulted in increased organ sizes (Doebley et al. 2006; Purugganan and Fuller 2009). The roots of cultivated manioc have greater amounts and larger starch granules, but not larger cell sizes, than the wild relatives (Lebot 2009; An et al. 2016). However, the cessation of cell division and expansion may play an important role during manioc response to drought stress (Alves 2002; Alves and Setter 2004). Another outlier SNP (*Pst_7007*) was in a gene that may act in responses to salt, drought, and cold stresses. Responses to abiotic stresses were important crop adaptations for the ecological shifts from the wild to the cultivation environment and when they started to be dispersed (McKey et al. 2012; Meyer and Purugganan 2013). In the specific case of manioc, abiotic resistance is relevant to crop adaptability to marginal areas and might be key for manioc adaption to harsher future climates (Jarvis et al. 2012).

We recognize that the type of genomic library and the limited sampling number of some groups of varieties may have introduced bias in the analyses (Nielsen and Signorovitch 2003; Arnold et al. 2013). Although the groups of manioc varieties are not true populations, some aspects, such as the possibility of crossings and incorporation of sexual plants into clonally propagated varieties, make these groups biologically meaningful. Moreover, genetic approaches provided interesting results even when using generic groups of manioc varieties (Bradbury et al. 2013; Alves-Pereira et al. 2020) and limited sample sizes in other study systems (Talavera et al. 2019; Kates et al. 2021). The selective signatures discussed above should be regarded as initial hypotheses and further genome-wide association studies and quantitative trait loci mapping are required to confirm their biological significance (Barrett and Hoekstra 2011; Meyer and Purugganan 2013). Nonetheless, this new information may guide future bottom-up approaches to characterize the genomic changes and their associated phenotypic effects (Ross-Ibarra et al. 2007) relevant to manioc evolution under domestication and to assist breeding strategies, contributing to our understanding about adaptive genes in crops.

### Genome-wide diversity

The genetic diversity of varieties from different biomes (*H*_*E*_ ranging from 0.285 to 0.312) was similar to that observed in previous studies and suggests that the gene bank conserves high levels of genetic diversity, despite its limited size. Within Embrapa Brazilian manioc germplasm bank, Albuquerque et al. (2018) reported an overall *H*_*E*_ = 0.29 (biallelic SNPs) and Ogbonna et al. (Ogbonna et al. 2021a) found *H*_*O*_ ranging from 0.26 to 0.39 across genetic groups. Similar results were observed for the gene banks of the International Institute of Tropical Agriculture (IITA, *H*_*E*_= 0.334) and the International Center for Tropical Agriculture (CIAT, *H*_*E*_ = 0.341) (Ferguson et al. 2019). This high genetic diversity is explained by the complex evolutionary dynamics of the crop under traditional cultivation, that results in varieties consisting of a predominant clone plus individuals morphologically similar, but genetically distinct (Peroni et al. 2007). Although smallholder farmers propagate manioc exclusively by stem cuttings, crossings between different varieties may occur producing sexual seeds that become part of the soil seed bank and may sprout amid clonally propagated plants (Duputié et al. 2009; McKey et al. 2010). After harvesting, the farmers may incorporate stem cuttings of these sexual plants into their clonal stocks leading to the amplification and maintenance of genetic diversity in the local scale (Martins 2007). The high genetic diversity observed in the different biomes may be explained by the widespread occurrence of the incorporation of sexual plants to clonal varieties, as well as by the selection for distinct human preferences under diverse ecological and cultural contexts (Clement et al. 2010; Gepts 2014). Genetic evidence for different human preferences in distinct ecogeographic contexts has already been reported in different regions of South America for manioc (Peña-Venegas et al. 2014; Mühlen et al. 2019) and other crops (Clement et al. 2017; Moreira et al. 2017).

The crop’s reproductive biology and other traditional farming practices may explain the low to moderate genetic divergence among biomes (*F*_*ST*_ ranging from -0.025 to 0.065) and the high admixture observed in clustering analyses (Figure 3). Exchange networks of manioc varieties have been reported at local and ample geographic scales (Heckler and Zent 2008; Rabbi et al. 2015). These exchange networks facilitate geneflow between distinct varieties at local and broader geographical scales. Genetic admixture among varieties from distinct geographical locations are commonly associated with extensive exchange networks of manioc and other crops (Delêtre et al. 2011; Mutegi et al. 2011; Roullier et al. 2011). This low overall genetic divergence may also reflect the initial common dispersal of landraces from their domestication center in Amazonia (Alves-Pereira et al. 2018; Mühlen et al. 2019). The Amazonian varieties were somewhat more divergent in relation to the other biomes (Figure 3D, Table 4), possibly because almost all bitter varieties were from this region. The influence of ecogeographic variation in the distribution of genetic diversity in manioc is variable (Siqueira et al. 2009; Mühlen et al. 2019; Ogbonna et al. 2021a), but the existence of divergent varieties from different biomes may also reflect the selection under distinct ecological and cultural contexts. Nonetheless, the genetic divergence between bitter and sweet manioc seems to be higher in Amazonia than outside this region (Bradbury et al. 2013; Alves-Pereira et al. 2020). It is possible that divergent selective pressures for bitterness or sweetness are more relaxed outside Amazonia, because the crop’s dispersals were not always accompanied by cultural appropriation (McKey and Delêtre 2017). Knowledge about adequate processing to avoid intoxication after the consumption of bitter manioc landraces is essential to achieve food security where people rely on manioc cultivation (Burns et al. 2010).

The remarkable divergence between wild and cultivated manioc was expected given the long history of the crop diversification under human selection and cultivation (Olsen and Schaal 1999). This result may also reflect many founding events (Allaby et al. 2019) that accompanied the rapid spread of the crop across the Neotropics (Isendahl 2011) and the wide dispersal across the world. The observed genetic divergence was similar to our previous study (Alves-Pereira et al. 2020), but more geographically extensive sampling of wild populations would improve our understanding about the current population dynamics between wild and cultivated manioc. Recent genomic approaches evidenced introgressions from some wild relatives in the genome of cultivated manioc (Bredeson et al. 2016; Wolfe et al. 2019). Because the primary gene pool of manioc consists of 13 species (Allem 2002), introgressions may have been contributing to the extant genetic diversity of the crop. Therefore, genome-wide studies in cultivated manioc and different wild *Manihot* species could greatly contribute to our understanding about the evolution of the crop.

In this study, we observed high levels of genome-wide diversity in manioc varieties from different Brazilian biomes. It is noteworthy that the genetic diversity also included putative adaptive variation, which may be associated with the crop’s domestication and its cultivation in distinct environmental contexts with different human preferences. Some of the signatures of selection may be associated with resistance genes and agronomic traits of interest, which might have practical importance for breeding purposes. The varieties conserved in the gene bank can be used as sources for reintroductions into smallholder communities (Barbieri et al. 2014): the highly genetic divergent varieties are important resources for their specific regions of origin, while the admixed varieties may adapt well to cultivation in various locations. This study reinforces the importance of *ex situ* collections for the conservation of the crops’ genetic resources, although it is financially and technically challenging to maintain active gene banks (Esquinas-Alcázar 2005; Khoury et al. 2014). Our study also highlights the necessity of maintaining traditional practices of cultivation, since they are often associated with the management of a great diversity of other native crops (Peroni and Martins 2000; Peroni and Hanazaki 2002; Barbieri et al. 2014). In the context of a changing world, the characterization and conservation of agrobiodiversity is essential for the appropriate management of their genetic resources and ultimately for food security, especially of poor people.

## Data availability statement

Final SNP data uploaded as online Table S2. Sequence alignments (bam files) were deposited in NCBI SRA (accessions PRJNA748763 (library *Nsi*I+*Msp*I) and PRJNA748779 (library *Pst*I+*Mse*I)). These data will be fully available when the manuscript is accepted for publication with peer review.

## Conflicts of interest

The authors declare no conflicts of interests.

## Author contributions

AA-P, MIZ and APS designed the research. CRC, MIZ, APS, EAV and JBP got financial support. AA-P, MIZ and APS performed the experiments. AA-P and JPGV performed the analyses. AA-P wrote the initial manuscript and all the other co-authors contributed to its final form.

**Appendix 1.** Predicted effects and annotations of 21 SNP markers with putative selective signatures considering different groups of manioc (*Manihot esculenta*) samples. a: SNP identification as specified in the vcf file; b: chromosome names are coded according to the manioc genome *Manihot esculenta v6* (NCBI PRJNA234389) and the SNP positions are within parenthesis; c: asterisks indicate putative manioc resistance genes (PRGdb v.3.0); d: B indicates that the tests were performed considering the groups of varieties per biome, while G the groups of wild and cultivated manioc; e: mutational effect predicted for the primary transcript of the associated manioc genes. Amino acid substitutions are coded as [number of the residue] original amino acid > new amino acid; f: E-value = probability of hits by chance; g: Recovered from the UniProt data base (www.uniprot.org).

| SNP ID <sup>a</sup> | Genomic location <sup>b</sup> | Gene name <sup>c</sup> | Outlier tests <sup>d</sup> | Predicted effect <sup>e</sup> | Name of the protein with Blastp similarity | Identity (%) | E-value <sup>f</sup> | Swiss-Prot Accession | Functional description <sup>g</sup> | Keywords |
| --- | --- | --- | --- | --- | --- | --- | --- | --- | --- | --- |
| Pst_1761 (G > T) | NC_035163.1 (2920331) | LOC110610533 (Manes.03G036300)* | G (FLK, pcadapt, FST) | Missense variant [280] Ala > Ser | Disease resistance protein RPS2 | 100 | 0 | Q42484 | RPS2 is a disease resistance protein involved in the response to *Pseudomonas syringae* (Mackey et al. 2003). | Resistance to pathogens |
| Pst_6563 (C > T) | NC_035172.1 (1844961) | LOC110628374 (Manes.12G024600)* | G (FLK, pcadapt) | Missense variant [604] Thr > Ile | Disease resistance protein RPM1 | 100 | 0 | Q39214 | RPM1 is a disease resistance protein involved in the response to *Pseudomonas syringae* (Mackey et al. 2002). | Resistance to pathogens |
| Nsi_172 (A > C) | NC_035161.1 (22495119) | LOC110630587 (Manes.01G102500)* | B (FLK, FST) | Synonymous variant [736] Leu > Leu | Receptor-like protein kinase FERONIA | 100 | 0 | Q9SCZ4 | FERONIA is a receptor-like protein kinase involved in growth cessation of compatible pollen tubes ensuring reproductive isolation barriers (Escobar-Restrepo et al. 2007). Required for cell elongation during vegetative growth (Guo et al. 2009). Mediates sensitivity to powdery mildew (Kessler et al. 2010). | Fertilization, Cell elongation, Powdery mildew |
| Pst_4209 (A > G) | NC_035167.1 (4576622) | LOC110618886 (Manes.07G046400)* | G (FLK, pcadapt) | Missense variant [440] Val > Ala | Receptor-like protein kinase FERONIA | 100 | 0 | Q9SCZ4 | Idem. | Idem. |
| Pst_3333 (G > A) | NC_035165.1 (20713381) | LOC110616124 (Manes.05G145600)* | G (hapFLK, pcadapt) | 5' UTR variant | Receptor protein kinase CLAVATA1 | 100 | 0 | Q9SYQ8 | CLAVATA1 acts in a signal transduction pathway controlling the balance between meristem cell proliferation and differentiation (Bleckmann et al. 2010). | Development, Cell proliferation, Growth |
| Pst_1010 (C > T) | NC_035162.1 (1649693) | LOC110609084 (Manes.02G020500)* | G (FLK, pcadapt) | Missense variant [318] Ala > Thr | Mitogen-activated protein kinase kinase kinase YODA (YDA) | 99.4 | 0 | Q9CAD5 | YDA is involved in the first cell fate decisions in the zygote and the early embryo. It is involved in regulating coordinated local cell proliferation, which shapes the morphology of plant organs (Lukowitz et al. 2004). | Cell proliferation, Organ shape |
| Pst_6259 (C > G) | NC_035171.1 (14640434) | LOC110626307 (Manes.11G094000)* | B (hapFLK, FST) | Synonymous variant [604] Gly > Gly | Proline-rich receptor-like protein kinase PERK12 (PERK12) | 100 | 0 | Q9ZUE0 | PERK12 regulates the auxin-related MAX (More Axillary Growth) pathway during shoot branching (Hwang et al. 2010). | Branching |
| Nsi_4509 (A > T) | NC_035178.1 (777460) | LOC110606622 (Manes.18G008800)* | G (FLK, pcadapt) | Upstream gene variant | Serine/threonine-protein kinase (STY46) | 100 | 0 | F4JTP5 | SYT46 may be part of a cytosolic regulatory network involved in chloroplast protein import. Plays a role in chloroplast biogenesis and differentiation in cotyledons (Martin et al. 2006; Lamberti et al. 2011). | Chloroplast biogenesis and differentiation in cotyledons |
| Nsi_3361 (G > A) | NC_035173.1 (4723844) | LOC110630205 (Manes.13G043200) | B (FLK, FST) | Synonymous variant [148] Leu > Leu | 26S proteasome regulatory subunit 4 homolog A (RPT2A) | 96.4 | 0 | Q9SZD4 | RPT2A acts in degradation of ubiquitinated proteins and is required for the maintenance of postembryonic root and shoot meristems (Ueda et al. 2004). It is involved in the regulation of organ size (Sonoda et al. 2009), in the regulation of gametogenesis (Ueda et al. 2011), in the regulation of transcriptional and post-transcriptional gene silencing (Sako et al. 2012), tolerance to zinc deficiency (Sakamoto et al. 2011), and fungal resistance (Yao et al. 2012). | Development (root and shoot), Organ size, Defense, Fungal resistance, Gene silencing, Gametogenesis, Ubiquitin |
| Pst_4290<br>(G > C) | NC_035167.1<br>(19228818) | LOC110618716<br>(Manes.07G083000) | G (FLK,<br>pcadapt,<br>FST) | Intron variant | Ubiquitin C-terminal<br>hydrolase 12 (UBP12) | 85.6 | 0 | Q9FPT1 | UBP12 is involved in the processing of ubiquitinated proteins acting as positive regulator of root meristem development (An et al. 2018). | Root development, Ubiquitin |
| Nsi_2393<br>(C > A) | NC_035169.1<br>(5588020) | LOC110622644<br>(Manes.09G041200) | G (FLK,<br>pcadapt,<br>FST) | Intron variant | Ubiquitin C-terminal<br>hydrolase 12 (UBP12) | 87.3 | 0 | Q9FPT1 | Idem. | Root development, Ubiquitin |
| Pst_8996<br>(A > C) | NC_035176.1<br>(24796226) | LOC110603073<br>(Manes.16G091200) | G (FLK,<br>pcadapt,<br>FST) | Synonymous variant [3] Val > Val | Auxin response factor 19 (ARF19) | 83.2 | 0 | Q8RYC8 | Auxin response factors (ARFs) are transcriptional factors. ARF19 may regulate lateral root formation (Okushima et al. 2007). | Transcriptional factor, Root formation |
| Pst_5985<br>(G > C) | NC_035171.1<br>(262333) | LOC110626926<br>(Manes.11G002200) | G (FLK,<br>pcadapt,<br>FST) | 3' UTR variant | Glycosyltransferase-like KOBITO 1 (ELD1) | 77.1 | 0 | Q9C9Z9 | ELD1 is involved in the coordination between cell elongation and cellulose synthesis (Wang et al. 2015), in abscisic acid (ABA) and sugar responses essential for growth (Brocard-Gifford et al. 2004), maintaining normal organogenesis. It is involved with cell elongation and maintenance of root meristem identity (Lertpiriyapong and Sung 2003). Acts as negative factor in light inhibition of hypocotyl elongation (Wang et al. 2015). | Growth, Cell elongation |
| Pst_5701<br>(C > T) | NC_035170.1<br>(5198647) | LOC110624912<br>(Manes.10G051700) | G (FLK,<br>hapFLK,<br>pcadapt,<br>FST) | Intron variant | Pullulanase 1, chloroplastic (PU1) | 72.4 | 0 | Q8GTR4 | PU1 is involved in starch degradation and probably in starch synthesis (Wattebled et al. 2008). | Starch |
| Pst_7007<br>(C > T) | NC_035173.1<br>(1033679) | LOC110630343<br>(Manes.13G010500) | G (FLK,<br>pcadapt,<br>FST) | Missense_variant [932] Ser > Asn | Histidine kinase 3 (AHK3) | 72 | 0 | Q9C5U1 | AHK3 acts in the transmission of drought, salt, and cold stress signals (Tran et al. 2007; Jeon et al. 2010). It is involved in many developmental processes: meristem establishment in seedlings, seed germination, cell division, seed size, chlorophyll retention during leaf senescence, root repression and shoot promotion, and flower development (Higuchi et al. 2004; Riefler et al. 2006; Li et al. 2010). | Stress (salt, drought, cold), Development (flower, plant) |
| Pst_8971<br>(C > T) | NC_035176.1<br>(24448370) | LOC110604012<br>(Manes.16G087200) | G (FLK,<br>pcadapt) | Intron variant | 3beta-hydroxysteroid-dehydrogenase/decarboxylase isoform 2 (3BETAHSD/D2) | 71.9 | 0 | Q67ZE1 | 3BETAHSD/D2 is involved in the regulation of inflorescence internodes and leaf growth (Kim et al. 2012). | Growth (inflorescence, leaf) |
| Pst_29 (T > C) | NC_035161.1<br>(1823645) | LOC110613144<br>(Manes.01G010300) | B (hapFLK,<br>FST) | Synonymous variant [132] Val > Val | Thiocyanate methyltransferase 1 (TMT1) | 60.2 | $8.2 \times 10^{-70}$ | Q93V78 | TMT1 is a methyltransferase possibly involved in glucosinolate metabolism after tissue damage (Attieh et al. 2000). | Cyanide, Defense |
| Nsi_2289<br>(A > G) | NC_035168.1<br>(30311331) | LOC110620226<br>(Manes.08G136900) | G (hapFLK,<br>pcadapt) | Intron variant | GRF1-interacting factor 1 (GIF1) | 62.5 | $1.7 \times 10^{-62}$ | Q8L8A5 | GIF1 is a transcription coactivator involved in the regulation of cell expansion in leaves and cotyledons (Fujikura et al. 2009), acting in the development of appropriate leaf size and shape. It also mediates cotyledon identity (Kanei et al. 2012). | Development (leaf, cotyledon) |
| Nsi_3540<br>(C > A) | NC_035174.1<br>(731020) | LOC110599603<br>(Manes.14G007100) | G (FLK,<br>hapFLK,<br>pcadapt,<br>FST) | Missense variant [147] Ser > Arg | FT-interacting protein 3 (FTIP3) | 60.2 | 0 | Q9M2R0 | FTIP3 is required for the proliferation and differentiation of shoot stem cells in the shoot apical meristem (Liu et al. 2018). | Development of shoot |
| Pst_9853<br>(A > G) | NC_035178.1<br>(7429534) | LOC110606992<br>(Manes.18G086300) | G (FLK,<br>pcadapt) | Intron variant | Transcriptional corepressor SEUSS (SEU) | 62.9 | 0 | Q8W234 | SEU is involved in the transcriptional repression of the C class floral homeotic gene AGAMOUS during flower development (Sridhar et al. 2004), and in the regulation of petal shape and ovule development (Bao et al. 2010). | Flowering, Transcription regulation |
| Pst_9959<br>(A > G) | NC_035178.1<br>(11748439) | LOC110607096<br>(Manes.18G120200) | B (FLK,<br>hapFLK) | Missense variant [81] Asp > Gly | Agamous-like MADS-box protein AGL12 (AGL12) | 60.4 | $2.8 \times 10^{-78}$ | Q38841 | AGL12 may be a transcription activator involved in the regulation of root development by controlling cell proliferation. May act as promoter of flowering transition (Tapia-López et al. 2008). | Root, Flowering, Transcription regulation |

## Acknowledgments

This study was supported by various grants, including São Paulo Research Foundation (FAPESP 2018/00036-9, 2013/00003-0 and 2012/08307-5), and Conselho Nacional de Desenvolvimento CientÍfico e Tecnológico (CNPq, CT-Amazônia 575588/08-0). We thank Aline Moraes and Danilo Sforza for laboratory assistance. AA-P thanks FAPESP for a post-doctoral scholarship (2018/00036-9). APS thanks CNPq for a Research Fellowship (312777/2018–3) and Coordenação de Aperfeiçoamento de Pessoal de NÍvel Superior (CAPES - Computational Biology Program, 88882.160095/2013-01).

## Notes

### Competing Interest Statement

The authors have declared no competing interest.

## References

Ahrens CW, Rymer PD, Stow A, Bragg J, Dillon S, Umbers KDL, Dudaniec RY (2018) The search for loci under selection: trends, biases and progress. Mol Ecol 27:1342–1356. https://doi.org/10.1111/mec.14549

Albuquerque HYG, Carmo CD do, Brito AC, Oliveira EJ de (2018) Genetic diversity of Manihot esculenta Crantz germplasm based on single-nucleotide polymorphism markers. Ann Appl Biol 173:271–284. https://doi.org/10.1111/aab.12460

Alexa A, Rahnenführer J (2021) TopGO: Enrichment analysis for Gene Ontology. R package version 2.44.0.

Allaby RG, Ware RL, Kistler L (2019) A re-evaluation of the domestication bottleneck from archaeogenomic evidence. Evol Appl 12:29–37. https://doi.org/10.1111/eva.12680

Allem AC (2002) The origins and taxonomy of cassava. In: Hillocks RJ, Thresh JM, Bellotti A (eds) Cassava: Biology, Production and Utilization. CABI, Wallingford, UK, pp 1–16

Allem AC (1994) The origin of Manihot esculenta Crantz (Euphorbiaceae). Genet Resour Crop Evol 41:133–150. https://doi.org/10.1007/BF00051630

Allendorf FW (2017) Genetics and the conservation of natural populations: allozymes to genomes. Mol Ecol 26:420–430. https://doi.org/10.1111/mec.13948

Alves-Pereira A, Clement CR, Picanço-Rodrigues D, Veasey EA, Dequigiovanni G, Ramos SLF, Pinheiro JB, de Souza AP, Zucchi MI (2020) A population genomics appraisal suggests independent dispersals for bitter and sweet manioc in Brazilian Amazonia. Evol Appl 13:342–361. https://doi.org/10.1111/eva.12873

Alves-Pereira A, Clement CR, Picanço-Rodrigues D, Veasey EA, Dequigiovanni G, Ramos SLF, Pinheiro JB, Zucchi MI (2018) Patterns of nuclear and chloroplast genetic diversity and structure of manioc along major Brazilian Amazonian rivers. Ann Bot 121:625–639. https://doi.org/10.1093/aob/mcx190

Alves AAC (2002) Cassava botany and physiology. In: Hillocks RJ, Thresh JM, Bellotti A (eds) Cassava: Biology, Production and Utilization. CABI, Wallingford, UK, pp 67–89

Alves AAC, Setter TL (2004) Response of cassava leaf area expansion to water deficit: Cell proliferation, cell expansion and delayed development. Ann Bot 94:605–613. https://doi.org/10.1093/aob/mch179

An F, Chen T, Stéphanie DMA, Li K, Li QX, Carvalho LJCB, Tomlins K, Li J, Gu B, Chen S (2016) Domestication syndrome is investigated by proteomic analysis between cultivated cassava (Manihot esculenta Crantz) and its wild relatives. PLoS One 11:e0152154. https://doi.org/10.1371/journal.pone.0152154

An Z, Liu Y, Ou Y, Li J, Zhang B, Sun D, Sun Y, Tang W (2018) Regulation of the stability of RGF1 receptor by the ubiquitin-specific proteases UBP12/UBP13 is critical for root meristem maintenance. Proc Natl Acad Sci USA 115:1123–1128. https://doi.org/10.1073/pnas.1714177115

Andrews A (2010) FastQC: a quality control tool for high throughput sequence data [Online]. http://www.bioinformatics.babraham.ac.uk/projects/fastqc/

Arnold B, Corbett-Detig RB, Hartl D, Bomblies K (2013) RADseq underestimates diversity and introduces genealogical biases due to nonrandom haplotype sampling. Mol Ecol 22:3179– 3190. https://doi.org/10.1111/mec.12276

Attieh J, Sparace SA, Saini HS (2000) Purification and properties of multiple isoforms of a novel thiol methyltransferase involved in the production of volatile sulfur compounds from Brassica oleracea. Arch Biochem Biophys 380:257–266. https://doi.org/10.1006/abbi.2000.1896

Bao F, Azhakanandam S, Franks RG (2010) SEUSS and SEUSS-LIKE transcriptional adaptors regulate floral and embryonic development in Arabidopsis. Plant Physiol 152:821–836. https://doi.org/10.1104/pp.109.146183

Barbieri RL, Gomes JCC, Alercia A, Padulosi S (2014) Agricultural biodiversity in southern Brazil: Integrating efforts for conservation and use of neglected and underutilized species. Sustainability 6:741–757. https://doi.org/10.3390/su6020741

Barrett RDH, Hoekstra HE (2011) Molecular spandrels: tests of adaptation at the genetic level. Nat Rev Genet 12:767–780. https://doi.org/10.1038/nrg3015

Blake JA (2013) Ten quick tips for using the Gene Ontology. PLoS Comput Biol 9:e1003343. https://doi.org/10.1093/nar/gks1050.PLOS

Bleckmann A, Weidtkamp-Peters S, Seidel CAM, Simon R (2010) Stem cell signaling in Arabidopsis requires CRN to localize CLV2 to the plasma membrane. Plant Physiol 152:166–176. https://doi.org/10.1104/pp.109.149930

Bonhomme M, Chevalet C, Servin B, Boitard S, Abdallah J, Blott S, SanCristobal M (2010) Detecting selection in population trees: The Lewontin and Krakauer test extended. Genetics 186:241–262. https://doi.org/10.1534/genetics.110.117275

Bradbury EJ, Duputié A, Delêtre M, Roullier C, Narváez-Trujillo A, Manu-Aduening JA, Emshwiller E, McKey D (2013) Geographic differences in patterns of genetic differentiation among bitter and sweet manioc (Manihot esculenta subsp. esculenta; Euphorbiaceae). Am J Bot 100:857–866. https://doi.org/10.3732/ajb.1200482

Bredeson J V., Lyons JB, Prochnik SE, Wu GA, Ha CM, et al (2016) Sequencing wild and cultivated cassava and related species reveals extensive interspecific hybridization and genetic diversity. Nat Biotechnol 34:562–570. https://doi.org/10.1038/nbt.3535

Brocard-Gifford I, Lynch TJ, Garcia ME, Malhotra B, Finkelstein RR (2004) The Arabidopsis thaliana Abscisic Acid-Insensitive8 locus encodes a novel protein mediating abscisic acid and sugar responses essential for growth. Plant Cell 16:406–421. https://doi.org/10.1105/tpc.018077

Brown CH, Clement CR, Epps P, Luedeling E, Wichmann S (2013) The Paleobiolinguistics of domesticated manioc (Manihot esculenta). Ethnobiol Lett 4:61–70

Brown TA (2019) Is the domestication bottleneck a myth? Nat Plants 5:337–338. https://doi.org/10.1038/s41477-019-0404-1

Burns AE, Gleadow R, Cliff J, Zacarias A, Cavagnaro T (2010) Cassava: the drought, war and famine crop in a changing world. Sustainability 2:3572–3607. https://doi.org/10.3390/su2113572

Camacho C, Coulouris G, Avagyan V, Ma N, Papadopoulos J, Bealer K, Madden TL (2009) BLAST+: architecture and applications. BMC Bioinformatics 10:421. https://doi.org/10.1186/1471-2105-10-421

Castañeda-Álvarez NP, Khoury CK, Achicanoy HA, Bernau V, Dempewolf H, et al (2016) Global conservation priorities for crop wild relatives. Nat Plants 2:16022. https://doi.org/10.1038/NPLANTS.2016.22

Catchen J, Hohenlohe PA, Bassham S, Amores A, Cresko WA (2013) Stacks: An analysis tool set for population genomics. Mol Ecol 22:3124–3140. https://doi.org/10.1111/mec.12354

Ceballos H, Sánchez T, Morante N, Fregene M, Dufour D, Smith AM, Denyer K, Pérez JC, Calle F, Mestres C (2007) Discovery of an amylose-free starch mutant in cassava (Manihot esculenta Crantz). J Agric Food Chem 55:7469–7476. https://doi.org/10.1021/jf070633y

Champion A, Kreis M, Mockaitis K, Picaud A, Henry Y (2004) Arabidopsis kinome: after the casting. Funct Integr Genomics 4:163–187. https://doi.org/10.1007/s10142-003-0096-4

Cingolani P, Platts A, Wang LL, Coon M, Nguyen T, Wang L, Land SJ, Ruden DM, Lu X (2012) A program for annotating and predicting the effects of single nucleotide polyorphisms, SnpEff: SNPs in the genome of Drosophila melanogaster strain w1118; iso-2; iso-3. Fly (Austin) 6:1–13. https://doi.org/10.4161/fly.19695

Clement CR, Cristo-Araújo M de Coppensd ’Eeckenbrugge G, Reis VM dos, Lehnebach R, Picanço-Rodrigues D (2017) Origin and dispersal of domesticated peach palm. Front Ecol Evol 5:148. https://doi.org/10.3389/fevo.2017.00148

Clement CR, de Cristo-Araújo M, Coppensd’Eeckenbrugge G, Alves Pereira A, Picanço-Rodrigues D (2010) Origin and domestication of native Amazonian crops. Diversity 2:72– 106. https://doi.org/10.3390/d2010072

Coomes OT (2010) Of stakes, stems, and cuttings: The importance of local seed systems in traditional Amazonian societies. Prof Geogr 62:323–334. https://doi.org/10.1080/00330124.2010.483628

Danecek P, Auton A, Abecasis G, Albers CA, Banks E, DePristo MA, Handsaker RE, Lunter G, Marth GT, Sherry ST, McVean G, Durbin R, Group 1000 Genomes Project Analysis (2011) The variant call format and VCFtools. Bioinformatics 27:2156–2158. https://doi.org/10.1093/bioinformatics/btr330

Delêtre M, McKey D, Hodkinson TR (2011) Marriage exchanges, seed exchanges, and the dynamics of manioc diversity. Proc Natl Acad Sci USA 108:18249–18254. https://doi.org/10.1073/pnas.1106259108

Doebley JF, Gaut BS, Smith BD (2006) The molecular genetics of crop domestication. Cell 127:1309–1321. https://doi.org/10.1016.j.cell.2006.12.006

Doyle JJ, Doyle JL (1987) A rapid DNA isolation procedure for small quantities of fresh leaf tissue. Phytochem. Bull. 19:11–15

Duputié A, Massol F, David P, Haxaire C, McKey D (2009) Traditional Amerindian cultivators combine directional and ideotypic selection for sustainable management of cassava genetic diversity. J Evol Biol 22:1317–1325. https://doi.org/10.1111/j.1420-9101.2009.01749.x

Dyer GA, González C, Lopera DC (2011) Informal “seed” systems and the management of gene flow in traditional agroecosystems: The case of cassava in Cauca, Colombia. PLoS One 6:e29067. https://doi.org/10.1371/journal.pone.0029067

Elias M, Lenoir H, McKey D (2007) Propagule quantity and quality in traditional Makushi farming of cassava (Manihot esculenta): A case study for understanding domestication and evolution of vegetatively propagated crops. Genet Resour Crop Evol 54:99–115. https://doi.org/10.1007/s10722-005-2022-1

Elias M, McKey D (2000) The unmanaged reproductive ecology of domesticated plants in traditional agroecosystems: An example involving cassava and a call for data. Acta Oecologica 21:223–230. https://doi.org/10.1016/S1146-609X(00)00053-9

Elias M, Penet L, Vindry P, McKey D, Panaud O, Robert T (2001) Unmanaged sexual reproduction and the dynamics of genetic diversity of a vegetatively propagated crop plant, cassava (Manihot esculenta Crantz), in a traditional farming system. Mol Ecol 10:1895–1907. https://doi.org/10.1046/j.0962-1083.2001.01331.x

Ellen R, Soselisa HL (2012) A comparative study of the socio-ecological concomitants of cassava (Manihot esculenta Crantz) diversity, local knowledge and management in Eastern Indonesia. Ethnobot Res Appl 10:15–35. https://doi.org/10.17348/era.10.0.015-035

Escobar-Restrepo J-M, Huck N, Kessler S, Gagliardini V, Gheyselinck J, Yang W-C, Grossniklaus U (2007) The FERONIA receptor-like kinase mediates male-female interactions during pollen tube reception. Science (80-) 317:656–660. https://doi.org/10.1126/science.1143562

Esquinas-Alcázar J (2005) Protecting crop genetic diversity for food security: political, ethical and technical challenges. Nat Rev Genet 6:946–953. https://doi.org/10.1038/nrg1729

Excoffier L, Lischer HEL (2010) Arlequin suite ver 3.5: a new series of programs to perform population genetics analyses under Linux and Windows. Mol Ecol Resour 10:564–567. https://doi.org/10.1111/j.1755-0998.2010.02847.x

FAO (2010) The second report on the state of the world’s animal genetic resources for food and agriculture. FAO, Rome, Italy

FAO, IFAD, UNICEF, WFP, WHO (2021) The State of Food Security and Nutrition in the World 2021. Transforming food systems for food security, improved nutrition and affordable healthy diets for all. FAO, Rome, Italy

FAOSTAT (2019) Food and agriculture data. http://www.fao.org/faostat/en/#data/QC. Accessed 15 Jul 2021

Fariello MI, Boitard S, Naya H, SanCristobal M, Servin B (2013) Detecting signatures of selection through haplotype differentiation among hierarchically structured populations. Genetics 193:929–941. https://doi.org/10.1534/genetics.112.147231

Ferguson ME, Shah T, Kulakow P, Ceballos H (2019) A global overview of cassava genetic diversity. PLoS One 14:e0224763. https://doi.org/10.1371/journal.pone.0224763

Fernández-Llamazares Á, Lepofsky D, Lertzman K, Armstrong CG, Brondizio ES, et al (2021) Scientists’ warning to humanity on threats to indigenous and local knowledge systems. J Ethnobiol 41:144–169. https://doi.org/10.2993/0278-0771-41.2.144

Frichot E, François O (2015) LEA: An R package for landscape and ecological association studies. Methods Ecol Evol 6:925–929. https://doi.org/10.1111/2041-210X.12382

Frichot E, Mathieu F, Trouillon T, Bouchard G, François O (2014) Fast and efficient estimation of individual ancestry coefficients. Genetics 196:973–983. https://doi.org/10.1534/genetics.113.160572

Fujikura U, Horiguchi G, Ponce MR, Micol JL, Tsukaya H (2009) Coordination of cell proliferation and cell expansion mediated by ribosome-related processes in the leaves of Arabidopsis thaliana. Plant J 59:499–508. https://doi.org/10.1111/j.1365-313X.2009.03886.x

Gaillard MDP, Glauser G, Robert CAM, Turlings TCJ (2018) Fine-tuning the “plant domestication-reduced defense” hypothesis: specialist vs generalist herbivores. New Phytol 217:355–366. https://doi.org/10.1111/nph.14757

Gautier M, Klassmann A, Vitalis R (2017) rehh 2.0: a reimplementation of the R package rehh to detect positive selection from haplotype structure. Mol Ecol Resour 17:78–90. https://doi.org/10.1111/1755-0998.12634

Gepts P (2006) Plant genetic resources conservation and utilization: the accomplishments and future of a societal insurance policy. Crop Sci 46:2278–2292. https://doi.org/10.2135/cropsci2006.03.0169gas

Gepts P (2014) The contribution of genetic and genomic approaches to plant domestication studies. Curr Opin Plant Biol 18:51–59. https://doi.org/10.1016/j.pbi.2014.02.001

Godfray HCJ, Beddington JR, Crute IR, Haddad L, Lawrence D, Muir JF, Pretty J, Robinson S, Thomas SM, Toulmin C (2010) Food security: the challenge of feeding 9 billion people. Science (80-) 327:812–818. https://doi.org/10.1126/science.1185383

Guo H, Li L, Ye H, Yu X, Algreen A, Yin Y (2009) Three related receptor-like kinases are required for optimal cell elongation in Arabidopsis thaliana. Proc Natl Acad Sci USA 106:7648– 7653. https://doi.org/10.1073/pnas.0812346106

Halkier BA, Gershenzon J (2006) Biology and biochemistry of glucosinolates. Annu Rev Plant Biol 57:303–333. https://doi.org/10.1146/annurev.arplant.57.032905.105228

Hanks SK (2003) Genomic analysis of the eukaryotic protein kinase superfamily: a perspective. Genome Biol 4:111. https://doi.org/10.1186/gb-2003-4-5-111

Heckler S, Zent S (2008) Piaroa manioc varietals: Hyperdiversity or social currency? Hum Ecol 36:679–697. https://doi.org/10.1007/s10745-008-9193-2

Higuchi M, Pischke MS, Mähönen AP, Miyawaki K, Hashimoto Y, Seki M, Kobayashi M, Shinozaki K, Kato T, Tabata S, Helariutta Y, Sussman MR, Kakimoto T (2004) In planta functions of the Arabidopsis cytokinin receptor family. Proc Natl Acad Sci USA 101:8821– 8826. https://doi.org/10.1073/pnas.0402887101

Hillocks RJ., Wydra K (2002) Bacterial, fungal and nematode diseases. In: Hillocks RJ, Thresh JM, Bellotti A (eds) Cassava: Biology, Production and Utilization. CABI, Wallingford, UK, pp 261–280

Hoban S, Kelley JL, Lotterhos KE, Antolin MF, Bradburd G, Lowry DB, Poss ML, Reed LK, Storfer A, Whitlock MC (2016) Finding the genomic basis of local adaptation: Pitfalls, practical solutions, and future directions. Am Nat 188:379–397. https://doi.org/10.1086/688018

Howeler R, Lutaladio N, Thomas G (2013) Save and grow: cassava. A guide to sustainable production intensification. Food and Agriculture Organization of the United Nations, Rome

Hwang I, Kim SY, Kim CS, Park Y, Tripathi GR, Kim SK, Cheong H (2010) Over-expression of the IGI1 leading to altered shoot-branching development related to MAX pathway in Arabidopsis. Plant Mol Biol 73:629–641. https://doi.org/10.1007/s11103-010-9645-0

Isendahl C (2011) The domestication and early spread of manioc (Manihot esculenta Crantz): a brief synthesis. Lat Am Antiq 22:452–468. https://doi.org/10.7183/1045-6635.22.4.452

Jarvis A, Ramirez-Villegas J, Campo BVH, Navarro-Racines C (2012) Is cassava the answer to African climate change adaptation? Trop Plant Biol 5:9–29. https://doi.org/10.1007/s12042-012-9096-7

Jennings DL, Iglesias C (2002) Breeding for crop improvement. In: Hillocks RJ, Thresh JM, Bellotti A (eds) Cassava: Biology, Production and Utilization. CABI, Wallingford, UK, pp 149–166

Jeon J, Kim NY, Kim S, Kang NY, Novák O, Ku SJ, Cho C, Lee DJ, Lee EJ, Strnad M, Kim J (2010) A subset of cytokinin two-component signaling system plays a role in cold temperature stress response in Arabidopsis. J Biol Chem 285:23371–23386. https://doi.org/10.1074/jbc.M109.096644

Jombart T, Ahmed I (2011) Genetics and population analysis. Adegenet 1.3-1: new tools for the analysis of genome-wide SNP data. Bioinformatics 27:3070–3071. https://doi.org/10.1093/bioinformatics/btr521

Jombart T, Devillard S, Balloux F (2010) Discriminant analysis of principal components: a new method for the analysis of genetically structured populations. BMC Genet 11:94. https://doi.org/10.1186/1471-2156-11-94

Kanei M, Horiguchi G, Tsukaya H (2012) Stable establishment of cotyledon identity during embryogenesis in Arabidopsis by ANGUSTIFOLIA3 and HANABA TARANU. Development 139:2436–2446. https://doi.org/10.1242/dev.081547

Kates HR, Anido FL, Sánchez-de la Vega G, Eguiarte LE, Soltis PS, Soltis DE (2021) Targeted sequencing suggests wild-crop gene flow is central to different genetic consequences of two independent pumpkin domestications. Front Ecol Evol 9:618380. https://doi.org/10.3389/fevo.2021.618380

Keenan K, McGinnity P, Cross TF, Crozier WW, Prodöhl PA (2013) DiveRsity: an R package for the estimation and exploration of population genetics parameters and their associated errors. Methods Ecol Evol 4:782–788. https://doi.org/10.1111/2041-210X.12067

Kessler SA, Shimosato-Asano H, Keinath NF, Wuest SE, Ingram G, Panstruga R, Grossniklaus U (2010) Conserved molecular components for pollen tube reception and fungal invasion. Science (80-) 330:968–971. https://doi.org/10.1126/science.1195211

Khoury CK, Bjorkman AD, Dempewolf H, Ramirez-Villegas J, Guarino L, Jarvis A, Rieseberg LH, Struik PC (2014) Increasing homogeneity in global food supplies and the implications for food security. Proc Natl Acad Sci USA 111:4001–4006. https://doi.org/10.1073/pnas.1313490111

Kim B, Kim G, Fujioka S, Takatsuto S, Choe S (2012) Overexpression of 3β-hydroxysteroid dehydrogenases/C-4 decarboxylases causes growth defects possibly due to abnormal auxin transport in Arabidopsis. Mol Cells 34:77–84. https://doi.org/10.1007/s10059-012-0102-6

Kuon JE, Qi W, Schläpfer P, Hirsch-Hoffmann M, Von Bieberstein PR, Patrignani A, Poveda L, Grob S, Keller M, Shimizu-Inatsugi R, Grossniklaus U, Vanderschuren H, Gruissem W (2019) Haplotype-resolved genomes of geminivirus-resistant and geminivirus-susceptible African cassava cultivars. BMC Biol 17:1–15. https://doi.org/10.1186/s12915-019-0697-6

Lamberti G, Gügel IL, Meurer J, Soll J, Schwenkert S (2011) The cytosolic kinases STY8, STY17, and STY46 are involved in chloroplast differentiation in Arabidopsis. Plant Physiol 157:70– 85. https://doi.org/10.1104/pp.111.182774

Langmead B, Salzberg SL (2012) Fast gapped-read alignment with Bowtie 2. Nat Methods 9:357–359. https://doi.org/10.1038/nmeth.1923

Lebot V (2009) Tropical root and tuber crops: cassava, sweet potato, yams and aroids. CABI, Wallingford

Lenser T, Theißen G (2013) Molecular mechanisms involved in convergent crop domestication. Trends Plant Sci 18:704–714. https://doi.org/10.1016/j.tplants.2013.08.007

Léotard G, Duputié A, Kjellberg F, Douzery EJP, Debain C, de Granville JJ, McKey D (2009) Phylogeography and the origin of cassava: New insights from the northern rim of the Amazonian basin. Mol Phylogenet Evol 53:329–334. https://doi.org/10.1016/j.ympev.2009.05.003

Lertpiriyapong K, Sung ZR (2003) The elongation defective1 mutant of Arabidopsis is impaired in the gene encoding a serine-rich secreted protein. Plant Mol Biol 53:581–595. https://doi.org/10.1023/B:PLAN.0000019067.05185.d6

Li H (2011) A statistical framework for SNP calling, mutation discovery, association mapping and population genetical parameter estimation from sequencing data. Bioinformatics 27:2987–2993. https://doi.org/10.1093/bioinformatics/btr509

Li H, Handsaker B, Wysoker A, Fennell T, Ruan J, Homer N, Marth G, Abecasis G, Durbin R, Subgroup 1000 Genome Project Data Processing (2009) The Sequence Alignment/Map format and SAMtools. Bioinformatics 25:2078–2079. https://doi.org/10.1093/bioinformatics/btp352

Li XG, Su YH, Zhao XY, Li W, Gao XQ, Zhang XS (2010) Cytokinin overproduction-caused alteration of flower development is partially mediated by CUC2 and CUC3 in Arabidopsis. Gene 450:109–120. https://doi.org/10.1016/j.gene.2009.11.003

Liu L, Li C, Song S, Teo ZWN, Shen L, Wang Y, Jackson D, Yu H (2018) FTIP-dependent STM trafficking regulates shoot meristem development in Arabidopsis. Cell Rep 23:1879–1890. https://doi.org/10.1016/j.celrep.2018.04.033

Lotterhos KE, Whitlock MC (2015) The relative power of genome scans to detect local adaptation depends on sampling design and statistical method. Mol Ecol 24:1031–1046. https://doi.org/10.1111/mec.13100

Lotterhos KE, Whitlock MC (2014) Evaluation of demographic history and neutral parameterization on the performance of FST outlier tests. Mol Ecol 23:2178–2192. https://doi.org/10.1111/mec.12725

Lukowitz W, Roeder A, Parmenter D, Somerville C (2004) A MAPKK kinase gene regulates extra-embryonic cell fate in Arabidopsis. Cell 116:109–119. https://doi.org/10.1016/S0092-8674(03)01067-5

Luu K, Bazin E, Blum MGB (2017) pcadapt: an R package to perform genome scans for selection based on principal component analysis. Mol Ecol Resour 17:67–77. https://doi.org/10.1111/1755-0998.12592

Maccaferri M, Harris NS, Twardziok SO, Pasam RK, Gundlach H, et al (2019) Durum wheat genome highlights past domestication signatures and future improvement targets. Nat Genet 51:885–895. https://doi.org/10.1038/s41588-019-0381-3

Mackey D, Belkhadir Y, Alonso JM, Ecker JR, Dangl JL (2003) Arabidopsis RIN4 is a target of the type III virulence effector AvrRpt2 and modulates RPS2-mediated resistance. Cell 112:379–389. https://doi.org/10.1016/S0092-8674(03)00040-0

Mackey D, Holt BF, Wiig A, Dangl JL (2002) RIN4 interacts with Pseudomonas syringae type III effector molecules and is required for RPM1-mediated resistance in Arabidopsis. Cell 108:743–754. https://doi.org/10.1016/S0092-8674(02)00661-X

Martin T, Sharma R, Sippel C, Waegemann K, Soll J, Vothknecht UC (2006) A protein kinase family in Arabidopsis phosphorylates chloroplast precursor proteins. J Biol Chem 281:40216–40223. https://doi.org/10.1074/jbc.M606580200

Martins PS (2007) Dinâmica evolutiva em roças de caboclos amazônicos. In: Veasey EA, Oliveira GCX, Pinheiro JB (eds) Scientific papers of Paulo Sodero Martins 1941-1997: a tribute. SBG, Ribeirão Preto, pp 217–228

Mba REC, Stephenson P, Edwards K, Melzer S, Nkumbira J, Gullberg U, Apel K, Gale M, Tohme J, Fregene M (2001) Simple sequence repeat (SSR) markers survey of the cassava (Manihot esculenta Crantz) genome: Towards an SSR-based molecular genetic map of cassava. Theor Appl Genet 102:21–31. https://doi.org/10.1007/s001220051614

McCouch S, Baute GJ, Bradeen J, Bramel P, Bretting PK, et al (2013) Feeding the future. Nature 499:23–24. https://doi.org/10.1038/499023a

McKey D, Beckerman S (1993) Chemical ecology, plant evolution and traditional manioc cultivation systems. In: Hladik CM, Linares OF, Pagezy H, Semple A, Hadley M (eds) Tropical forests, people and food. Biocultural interactions and applications to development. Parthenon Carnforth and UNESCO, Paris, pp 83–112

McKey D, Cavagnaro TR, Cliff J, Gleadow R (2010) Chemical ecology in coupled human and natural systems: people, manioc, multitrophic interactions and global change. Chemoecology 20:109–133. https://doi.org/10.1007/s00049-010-0047-1

McKey D, Delêtre M (2017) The emergence of cassava as a global crop. In: Hershey CH (ed) Achieving sustainable cultivation of cassava, volume 1. Burleigh Dodds Science Publishing, London, pp 3–32

McKey D, Elias M, Pujol B, Duputié A (2012) Ecological approaches to crop domestication. In: Gepts P, Flamula TR, Bettinger RL, Brush SB, Damania AB, McGuire PE, Qualset CO (eds) Biodiversity in Agriculture: Domestication, Evolution, and Sustainability. Cambridge University Press, Cambridge, pp 377–406

Ménard L, McKey D, Mühlen GS, Clair B, Rowe NP (2013) The evolutionary fate of phenotypic plasticity and functional traits under domestication in manioc: changes in stem biomechanics and the appearance of stem brittleness. PLoS One 8:e74727. https://doi.org/10.1371/journal.pone.0074727

Meng X, Zhang S (2013) MAPK cascades in plant disease resistance signaling. Annu Rev Phytopathol 51:245–266. https://doi.org/10.1146/annurev-phyto-082712-102314

Meyer RS, DuVal AE, Jensen HR (2012) Patterns and processes in crop domestication: an historical review and quantitative analysis of 203 global food crops. New Phytol 196:29– 48. https://doi.org/10.1111/j.1469-8137.2012.04253.x

Meyer RS, Purugganan MD (2013) Evolution of crop species: Genetics of domestication and diversification. Nat Rev Genet 14:840–852. https://doi.org/10.1038/nrg3605

Moreira PA, Aguirre-Dugua X, Mariac C, Zekraoui L, Couderc M, Rodrigues DP, Casas A, Clement CR, Vigouroux Y (2017) Diversity of treegourd (Crescentia cujete) suggests introduction and prehistoric dispersal routes into Amazonia. Front Ecol Evol 5:150. https://doi.org/10.3389/fevo.2017.00150

Morrell PL, Buckler ES, Ross-Ibarra J (2012) Crop genomics: Advances and applications. Nat Rev Genet 13:85–96. https://doi.org/10.1038/nrg3097

Mühlen GS, Alves-Pereira A, Carvalho CRL, Junqueira AB, Clement CR, Valle TL (2019) Genetic diversity and population structure show different patterns of diffusion for bitter and sweet manioc in Brazil. Genet Resour Crop Evol 66:1773–1790. https://doi.org/10.1007/s10722-019-00842-1

Mutegi E, Sagnard F, Semagn K, Deu M, Muraya M, Kanyenji B, de Villiers S, Kiambi D, Herselman L, Labuschagne M (2011) Genetic structure and relationships within and between cultivated and wild sorghum (Sorghum bicolor (L.) Moench) in Kenya as revealed by microsatellite markers. Theor Appl Genet 122:989–1004. https://doi.org/10.1007/s00122-010-1504-5

Nielsen R, Signorovitch J (2003) Correcting for ascertainment biases when analyzing SNP data: applications to the estimation of linkage disequilibrium. Theor Popul Biol 63:245–255. https://doi.org/10.1016/S0040-5809(03)00005-4

Ogbonna AC, Braatz de Andrade LR, Mueller LA, de Oliveira EJ, Bauchet GJ (2021a) Comprehensive genotyping of a Brazilian cassava (Manihot esculenta Crantz) germplasm bank: insights into diversification and domestication. Theor Appl Genet. https://doi.org/10.1007/s00122-021-03775-5

Ogbonna AC, Braatz de Andrade LR, Rabbi IY, Mueller LA, Jorge de Oliveira E, Bauchet GJ (2021b) Large-scale genome-wide association study, using historical data, identifies conserved genetic architecture of cyanogenic glucoside content in cassava (Manihot esculenta Crantz) root. Plant J 105:754–770. https://doi.org/10.1111/tpj.15071

Okushima Y, Fukaki H, Onoda M, Theologis A, Tasaka M (2007) ARF7 and ARF19 regulate lateral root formation via direct activation of LBD/ASL genes in Arabidopsis. Plant Cell 19:118–130. https://doi.org/10.1105/tpc.106.047761

Oliveira EJ de, Resende MDV de, Santos V da S, Ferreira CF, Oliveira GAF, Silva MS da, Oliveira LA de, Aguilar-Vildoso CI (2012) Genome-wide selection in cassava. Euphytica 187:263– 276. https://doi.org/10.1007/s10681-012-0722-0

Olsen KM (2004) SNPs, SSRs and inferences on cassava’s origin. Plant Mol Biol 56:517–526. https://doi.org/10.1007/s11103-004-5043-9

Olsen KM, Schaal BA (1999) Evidence on the origin of cassava: Phylogeography of Manihot esculenta. Proc Natl Acad Sci USA 96:5586–5591. https://doi.org/10.1073/pnas.96.10.5586

Olsen KM, Schaal BA (2001) Microsatellite variation in cassava (Manihot esculenta, Euphorbiaceae) and its wild relatives: Further evidence for a southern Amazonian origin of domestication. Am J Bot 88:131–142. https://doi.org/10.2307/2657133

Osuna-Cruz CM, Paytuvi-Gallart A, Di Donato A, Sundesha V, Andolfo G, Cigliano RA, Sanseverino W, Ercolano MR (2018) PRGdb 3.0: a comprehensive platform for prediction and analysis of plant disease resistance genes. Nucleic Acids Res 46:D1197–D1201. https://doi.org/10.1093/nar/gkx1119

Pankin A, Altmüller J, Becker C, von Korff M (2018) Targeted resequencing reveals genomic signatures of barley domestication. New Phytol 218:1247–1259. https://doi.org/10.1111/nph.15077

Paquette SR (2012) Useful functions for (batch) file conversion and data resampling in microsatellite datasets. https://cran.r-project.org/package=PopGenKit

Peña-Venegas CP, Stomph TJ, Verschoor G, Lopez-Lavalle LAB, Struik PC (2014) Differences in manioc diversity among five ethnic groups of the Colombian Amazon. Diversity 6:792– 826. https://doi.org/10.3390/d6040792

Peroni N, Hanazaki N (2002) Current and lost diversity of cultivated varieties, especially cassava, under swidden cultivation systems in the Brazilian Atlantic Forest. Agric Ecosyst Environ 92:171–183. https://doi.org/10.1016/S0167-8809(01)00298-5

Peroni N, Kageyama PY, Begossi A (2007) Molecular differentiation, diversity, and folk classification of “sweet” and “bitter” cassava (Manihot esculenta) in Caiçara and Caboclo management systems (Brazil). Genet Resour Crop Evol 54:1333–1349. https://doi.org/10.1007/s10722-006-9116-2

Peroni N, Martins PS (2000) Influência da dinâmica agrÍcola itinerante na geração de diversidade de etnovariedades cultivadas vegetativamente. Interciencia 25:22–29

Poland JA, Brown PJ, Sorrells ME, Jannink JL (2012) Development of high-density genetic maps for barley and wheat using a novel two-enzyme genotyping-by-sequencing approach. PLoS One 7:e32253. https://doi.org/10.1371/journal.pone.0032253

Pritchard JK, Stephens M, Donnelly P (2000) Inference of population structure using multilocus genotype data. Genetics 155:945–959. https://doi.org/10.1111/j.1471-8286.2007.01758.x

Prochnik S, Marri PR, Desany B, Rabinowicz PD, Kodira C, Mohiuddin M, Rodriguez F, Fauquet C, Tohme J, Harkins T, Rokhsar DS, Rounsley S (2012) The cassava genome: current progress, future directions. Trop Plant Biol 5:88–94. https://doi.org/10.1007/s12042-011-9088-z

Pujol B, David P, McKey D (2005a) Microevolution in agricultural environments: How a traditional Amerindian farming practice favours heterozygosity in cassava (Manihot esculenta Crantz, Euphorbiaceae). Ecol Lett 8:138–147. https://doi.org/10.1111/j.1461-0248.2004.00708.x

Pujol B, Mühlen G, Garwood N, Horoszowski Y, Douzery EJP, McKey D (2005b) Evolution under domestication: contrasting functional morphology of seedlings in domesticated cassava and its closest wild relatives. New Phytol 166:305–318. https://doi.org/10.1111/j.1469-8137.2004.01295.x

Purugganan MD, Fuller DQ (2009) The nature of selection during plant domestication. Nature 457:843–848. https://doi.org/10.1038/nature07895

Quinlan AR, Hall IM (2010) BEDTools: a flexible suite of utilities for comparing genomic features. Bioinformatics 26:841–842. https://doi.org/10.1093/bioinformatics/btq033

R Core Team (2018) A language and environment for statistical computing. In: R Found. Stat. Comput. Vienna, Austria. https://www.r-project.org/. Accessed 15 Jan 2018

Rabbi IY, Kulakow PA, Manu-Aduening JA, Dankyi AA, Asibuo JY, Parkes EY, Abdoulaye T, Girma G, Gedil MA, Ramu P, Reyes B, Maredia MK (2015) Tracking crop varieties using genotyping-by-sequencing markers: A case study using cassava (Manihot esculenta Crantz). BMC Genet 16:115. https://doi.org/10.1186/s12863-015-0273-1

Reynolds J, Weir BS, Cockerham CC (1983) Estimation of the coancestry coefficient: basis for a short-term genetic distance. Genetics 105:767–779

Riefler M, Novak O, Strnad M, Schmülling T (2006) Arabidopsis cytokinin receptors mutants reveal functions in shoot growth, leaf senescence, seed size, germination, root development, and cytokinin metabolism. Plant Cell 18:40–54. https://doi.org/10.1105/tpc.105.037796

Ross-Ibarra J, Morrell PL, Gaut BS (2007) Plant domestication, a unique opportunity to identify the genetic basis of adaptation. Proc Natl Acad Sci USA 104:8641–8648. https://doi.org/10.1073/pnas.0700643104

Roullier C, Rossel G, Tay D, McKey D, Lebot V (2011) Combining chloroplast and nuclear microsatellites to investigate origin and dispersal of New World sweet potato landraces. Mol Ecol 20:3963–3977. https://doi.org/10.1111/j.1365-294X.2011.05229.x

Sabeti PC, Varilly P, Fry B, Lohmueller J, Hostetter E, et al (2007) Genome-wide detection and characterization of positive selection in human populations. Nature 449:913–918. https://doi.org/10.1038/nature06250

Sakamoto T, Kamiya T, Sako K, Yamaguchi J, Yamagami M, Fujiwara T (2011) Arabidopsis thaliana 26S proteasome subunits RPT2a and RPT5a are crucial for zinc deficiency-tolerance. Biosci Biotechnol Biochem 75:561–567. https://doi.org/10.1271/bbb.100794

Sako K, Maki Y, Kanai T, Kato E, Maekawa S, Yasuda S, Sato T, Watahiki MK, Yamaguchi J (2012) Arabidopsis RPT2a, 19S proteasome subunit, regulates gene silencing via DNA methylation. PLoS One 7:e37086. https://doi.org/10.1371/journal.pone.0037086

Salick J, Cellinese N, Knapp S (1997) Indigenous diversity of cassava: Generation, maintenance, use and loss among the Amuesha, peruvian upper amazon. Econ Bot 51:6–19. https://doi.org/10.1007/BF02910400

Sambatti JBM, Martins PS, Ando A (2001) Folk taxonomy and evolutionary dynamics of cassava: a case study in Ubatuba, Brazil. Econ Bot 55:93–105

Sardos J, Duval MF, McKey D, Malapa R, Lebot V, Noyer JL (2008) Evolution of cassava (Manihot esculenta Crantz) after recent introduction into a South Pacific Island system: the contribution of sex to the diversification of a clonally propagated crop. Genome 51:912–921. https://doi.org/10.1139/g08-080

Scheet P, Stephens M (2006) A fast and flexible statistical model for large-scale population genotype data: Applications to inferring missing genotypes and haplotypic phase. Am J Hum Genet 78:629–644. https://doi.org/10.1086/502802

Siqueira MVBM, Queiroz-Silva JR, Bressan EA, Borges A, Pereira KJC, Pinto JG, Veasey EA (2009) Genetic characterization of cassava (Manihot esculenta) landraces in Brazil assessed with simple sequence repeats. Genet Mol Biol 32:104–110. https://doi.org/10.1590/S1415-47572009005000010

Sonoda Y, Sako K, Maki Y, Yamazaki N, Yamamoto H, Ikeda A, Yamaguchi J (2009) Regulation of leaf organ size by the Arabidopsis RPT2a 19S proteasome subunit. Plant J 60:68–78. https://doi.org/10.1111/j.1365-313X.2009.03932.x

Sridhar V V, Surendrarao A, Gonzalez D, Conlan RS, Liu Z (2004) Transcriptional repression of target genes by LEUNIG and SEUSS, two interacting regulatory proteins for Arabidopsis flower development. Proc Natl Acad Sci USA 101:11494–11499. https://doi.org/10.1073/pnas.0403055101

Talavera A, Soorni A, Bombarely A, Matas AJ, Hormaza JI (2019) Genome-wide SNP discovery and genomic characterization in avocado (Persea americana Mill.). Sci Rep 9:20137. https://doi.org/10.1038/s41598-019-56526-4

Tapia-López R, GarcÍa-Ponce B, Dubrovsky JG, Garay-Arroyo A, Pérez-RuÍz R V., Kim SH, Acevedo F, Pelaz S, Alvarez-Buylla ER (2008) An AGAMOUS-related MADS-box gene, XAL1 (AGL12), regulates root meristem cell proliferation and flowering transition in Arabidopsis. Plant Physiol 146:1182–1192. https://doi.org/10.1104/pp.107.108647

Tran LSP, Urao T, Qin F, Maruyama K, Kakimoto T, Shinozaki K, Yamaguchi-Shinozaki K (2007) Functional analysis of AHK1/ATHK1 and cytokinin receptor histidine kinases in response to abscisic acid, drought, and salt stress in Arabidopsis. Proc Natl Acad Sci USA 104:20623–20628. https://doi.org/10.1073/pnas.0706547105

Ueda M, Matsui K, Ishiguro S, Kato T, Tabata S, Kobayashi M, Seki M, Shinozaki K, Okada K (2011) Arabidopsis RPT2a encoding the 26S proteasome subunit is required for various aspects of root meristem maintenance, and regulates gametogenesis redundantly with its homolog, RPT2b. Plant Cell Physiol 52:1628–1640. https://doi.org/10.1093/pcp/pcr093

Ueda M, Matsui K, Ishiguro S, Sano R, Wada T, Paponov I, Palme K, Okada K (2004) The HALTED ROOT gene encoding the 26S proteasome subunit RPT2a is essential for the maintenance of Arabidopsis meristems. Development 131:2101–2111. https://doi.org/10.1242/dev.01096

United Nations (2015) Transforming our world: the 2030 agenda for sustainable development. In: United nations general assembly

Wang X, Jing Y, Zhang B, Zhou Y, Lin R (2015) Glycosyltransferase-like protein ABI8/ELD1/KOB1 promotes Arabidopsis hypocotyl elongation through regulating cellulose biosynthesis. Plant, Cell Environ 38:411–422. https://doi.org/10.1111/pce.12395

Wattebled F, Planchot V, Dong Y, Szydlowski N, Pontoire B, Devin A, Ball S, D’Hulst C (2008) Further evidence for the mandatory nature of polysaccharide debranching for the aggregation of semicrystalline starch and for overlapping functions of debranching enzymes in Arabidopsis leaves. Plant Physiol 148:1309–1323. https://doi.org/10.1104/pp.108.129379

Weir BS, Cockerham CC (1984) Estimating F-statistics for the analysis of population structure. Evolution (N Y) 38:1358–1370

Wolfe MD, Bauchet GJ, Chan AW, Lozano R, Ramu P, Egesi C, Kawuk R, Kulakow P, Rabbi I, Jannink J-L (2019) Historical introgressions from a wild relative of modern cassava improved important traits and may be under balancing selection. Genetics 213:1237– 1253. https://doi.org/10.1534/genetics.119.302757

Yao C, Wu Y, Nie H, Tang D (2012) RPN1a, a 26S proteasome subunit, is required for innate immunity in Arabidopsis. Plant J 71:1015–1028. https://doi.org/10.1111/j.1365-313X.2012.05048.x

Zohary D (2004) Unconscious selection and the evolution of domesticated plants. Econ Bot 58:5–10. https://doi.org/10.1663/0013-0001(2004)058[0005:usateo]2.0.co;2

