## Supplementary Figures for "Selective signatures and high genome-wide diversity in traditional Brazilian manioc (*Manihot esculenta* Crantz) varieties"

*Corresponding author:

### Supplementary Material

#### Supplementary Tables

**Table S1** – Passport data for the 78 manioc (*Manihot esculenta*) varieties conserved at the Luiz de Queiróz College of Agriculture gene bank, Piracicaba, São Paulo, Brazil, and information about 14 additional samples of cultivated and wild manioc sampled in a previous study (Alves-Pereira *et al*., 2020). (In a separate Excel file).

**Table S2** – Variant calling format (VCF) file containing information for the 11,782 SNP markers identified for the 92 manioc (*Manihot esculenta*) varieties, based on two double-digest genotyping-by-sequencing genomic libraries. (In a separate Excel file).

**Table S3** – Predicted effects of mutations associated with the 865 SNP markers putatively under selection. CHROM = chromosome identification, and POS = position of the SNP in the manioc genome coded according to *Manihot esculenta v6* (NCBI PRJNA234389). ID = identification of the SNP marker; REF = reference allele; ALT = alternative allele; ANN = annotation of the predicted effects. Predicted effects on more than one gene transcript or in downstream/upstream genes are specified in different ANN columns. (In a separate Excel file).

**Table S4** – Results from the enrichment analysis based on GO annotations of the 663 manioc predicted genes with SNPs putatively under selection. GO annotations and their descriptions are shown with their respective observed and expected counts, and associated p-values from Fisher's exact test (In a separate Excel file).

**Table S5 –** Summary of blastp results for 351 amino acid sequences from manioc genes (*Manihot esculenta v6*, NCBI PRJNA234389) with outlier SNPs. Similarities are shown for 45 amino acid sequences with identity > 90 % when compared to the predicted resistance genes for manioc (PRGdb v.3.0) and 306 amino acid sequences with identity > 60 % with proteins deposited in Swiss-Prot. A summary of putative gene functions is also present if available in UniProt (www.unipr ot.org). (In a separate Excel file).

**Table S6** – Comparison of genetic diversity and inbreeding estimates based on 10,917 neutral SNPs for the groups of 92 manioc varieties and the 78 varieties conserved in the gene bank (within brackets). Number of samples (N), Total number of alleles (A), percentage of polymorphic loci (%P), number of private alleles (PA), observed ($H_O$) and expected ($H_E$) heterozygosities, inbreeding coefficients (*f*), and 95 % confidence intervals (95%CI).

| Groups | N | *A* | *%P* | *PA* | $H_O$ (95%CI) | $H_E$ (95%CI) | *f* (95%CI) |
|---|---|---|---|---|---|---|---|
| Cultivated | 84 | 21,516 | 98.5 | 7054 | 0.324 (0.321; 0.328) | 0.315 (0.312; 0.318) | -0.030 (-0.071; 0.005) |
| | [78] | [21,487] | [98.4] | [7054] | [0.328 (0.324; 0.331)] | [0.313 (0.310; 0.316)] | [-0.046 (-0.089; -0.010)] |
| Bitter | 16 | 21,009 | 97.9 | 60 | 0.295 (0.291; 0.298) | 0.309 (0.306; 0.312) | 0.047 (-0.014; 0.093) |
| | [13] | [20,937] | [97.7] | [74] | [0.302 (0.297; 0.304)] | [0.309 (0.306; 0.312)] | [0.026 (-0.052; 0.081)] |
| Sweet | 43 | 21,435 | 98.2 | 43 | 0.322 (0.318; 0.326) | 0.308 (0.305; 0.310) | -0.047 (-0.105; -0.001) |
| | [40] | [21,370] | [97.9] | [38] | [0.325 (0.321; 0.328)] | [0.306 (0.303; 0.309)] | [-0.061 (-0.126; -0.011)] |
| Wild | 8 | 14,780 | 67.7 | 318 | 0.103 (0.098; 0.105) | 0.122 (0.118; 0.125) | 0.177 (-0.040; 0.354) |
| Biomes | | | | | | | |
| Amazonia | 22 | 21,240 | 98.8 | 153 | 0.293 (0.290; 0.296) | 0.312 (0.310; 0.315) | 0.062 (0.015; 0.097) |
| | [16] | [21,137] | [98.4] | [124] | [0.299 (0.295; 0.302)] | [0.312 (0.310; 0.315)] | [0.044 (-0.021; 0.093)] |
| Cerrado | 30 | 21,108 | 98.2 | 16 | 0.342 (0.337; 0.346) | 0.298 (0.295; 0.301) | -0.145 (-0.216; -0.090) |
| Atlantic Forest | 27 | 21,142 | 99.7 | 7 | 0.325 (0.321; 0.329) | 0.300 (0.297; 0.303) | -0.085 (-0.167; -0.026) |
| Pantanal | 5 | 19,604 | 92.7 | 0 | 0.349 (0.344; 0.354) | 0.285 (0.281; 0.288) | -0.226 (-0.492; -0.107) |

**Table S7** – Comparison of analyses of molecular variance based on 10,917 neutral SNPs, showing the genetic variation within and among hierarchical groups, considering all the 92 varieties and the 78 varieties conserved in the gene bank (within brackets). Degrees of freedom (Df).

| Source of variation | Df | Sum of squares | Variance components | Percentage of variance | ϕ-statistics |
|---|---|---|---|---|---|
| Between Wild and Cultivated | 1<br>[1] | 22,823.83<br>[23,022.38] | 729.25<br>[740.41] | 32.5<br>[32.6] | $\phi_{ST}$ = 0.32 (p < 0.001)<br>[0.326 (p < 0.001)] |
| Within Wild and Cultivated | 182<br>[170] | 276,088.28<br>[260,663.40] | 1,516.97<br>[1,533.31] | 67.5<br>[67.4] | |
| Total | 183<br>[171] | 298,912.11<br>[283,685.78] | 2,246.22<br>[2,273.72] | | |
| Among Bitter, Sweet, and Wild | 2<br>[2] | 26,517.04<br>[26,174.50] | 341.13<br>[375.32] | 19.1<br>[20.3] | $\phi_{ST}$ = 0.19 (p < 0.001)<br>[0.203 (p < 0.001)] |
| Within Bitter, Sweet, and Wild | 131<br>[119] | 189,459.88<br>[175,011.93] | 1,446.26<br>[1,470.68] | 80.9<br>[79.7] | |
| Total | 133<br>[121] | 215,976.92<br>[201,186.43] | 1,787.39<br>[1,846.01] | | |
| Among Biomes | 3<br>[3] | 11,073.90<br>[9,589.59] | 54.76<br>[45.18] | 3.4<br>[2.8] | $\phi_{ST}$ = 0.03 (p < 0.001)<br>[0.028 (p < 0.001)] |
| Within Biomes | 164<br>[152] | 254,820.13<br>[240,884.75] | 1,553.78<br>[1,584.77] | 96.6<br>[97.2] | |
| Total | 167<br>[155] | 265,894.03<br>[250,474.34] | 1,608.54<br>[1,629.95] | | |

| | | | | | |
|---|---|---|---|---|---|
| Between Biomes and Wild | 1 [1] | 22,823.83 [23,022.38] | 693.60 [710.26] | 31.1 [31.4] | $\phi_{ST}$ = 0.34 ($p < 0.001$) [0.335 ($p < 0.001$)] |
| Among groups within Biomes and Wild | 3 [3] | 11,073.90 [9,589.59] | 56.64 [47.46] | 2.5 [2.1] | $\phi_{SC}$ = 0.03 ($p < 0.001$) [0.003 ($p < 0.001$)] |
| Within groups | 179 [167] | 265,014.38 [251,073.81] | 1,480.53 [1,503.44] | 66.4 [66.5] | $\phi_{CT}$ = 0.31 ($p = 0.19$) [0.31 ($p = 0.20$)] |
| Total | 183 [171] | 298,912.11 [283,685.78] | 2,230.77 [2,261.15] | | |

**Table S8** – Comparison of pairwise estimates of genetic divergence (Weir & Cockerham's $F_{ST}$ (1984)) based on 10,917 neutral SNPs, considering all the 92 varieties and the 78 varieties conserved in the gene bank (within brackets). Asterisks indicate estimates significant at $p < 0.01$.

| Groups | Bitter | Sweet | | |
|---|---|---|---|---|
| Bitter | | | | |
| Sweet | 0.045* [0.044*] | | | |
| Wild | 0.357* [0.349*] | 0.344* [0.349*] | | |
| Biomes | Amazonia | Cerrado | Atlantic Forest | Pantanal |
| Amazonia | | | | |
| Cerrado | 0.065* [0.060*] | | | |
| Atlantic Forest | 0.045* [0.039*] | 0.017 | | |
| Pantanal | 0.044* [0.036] | -0.025 | -0.015 | |
| Wild | 0.342* [0.339*] | 0.365* | 0.362* | 0.433* |

**Supplementary Figures**

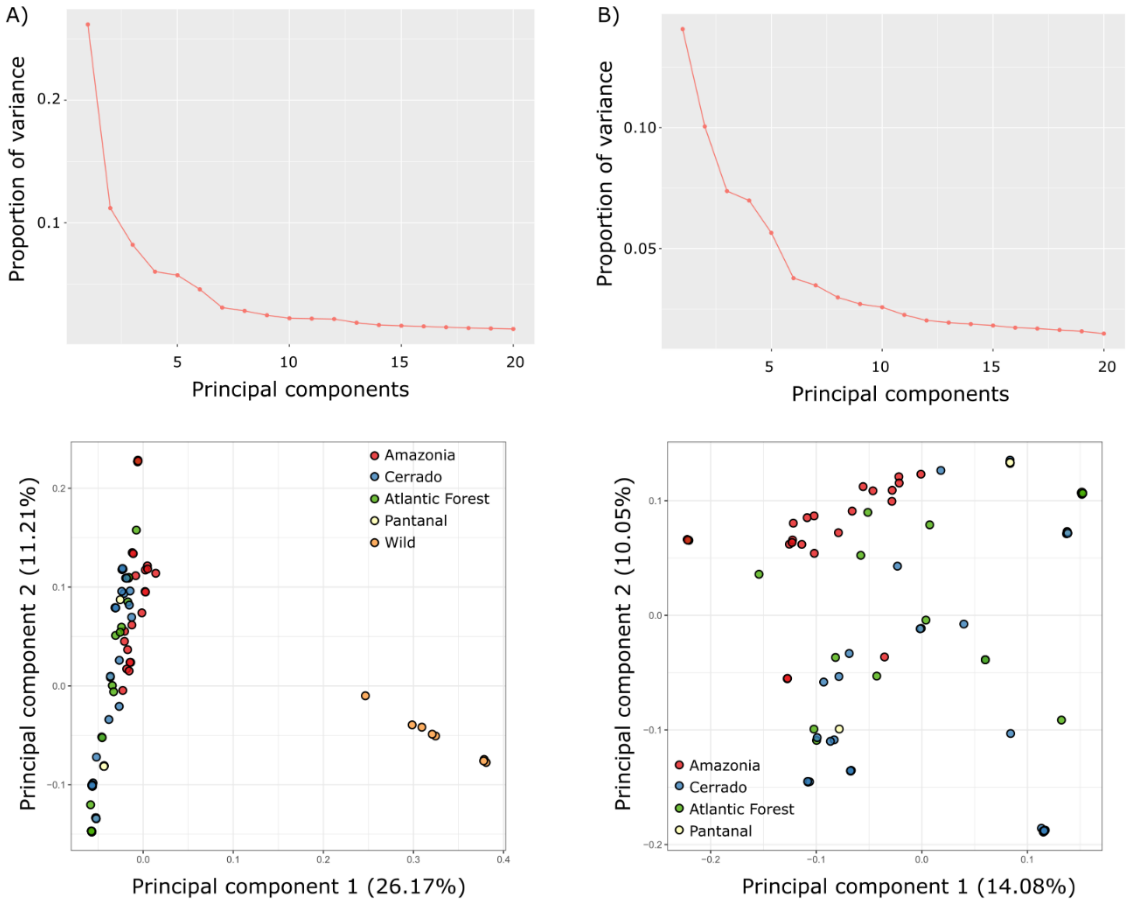

**Figure S1** – Pcadapt results for the detection of outlier SNPs considering A) the groups of wild versus cultivated manioc (*Manihot esculenta*, N = 92), and B) the groups of varieties per biomes (without wild samples, N = 84). The analyses were performed based on 11,782 SNP markers. Top: Scree plots of the proportion of explained variance in the principal component analysis (PCA) for the first K = 20 principal components. The number of components retained in the analyses followed Cattle's rule, choosing the point to the left where the curve inflects, and the addition of components does not substantially increase the amount of variance explained. We chose K = 2 for the analysis contrasting wild and cultivated manioc, and K = 5 for the analysis contrasting the biomes. Bottom: Scatter plots of the first two principal components showing the major genetic structure observed in the PCAs for the two groupings of samples.

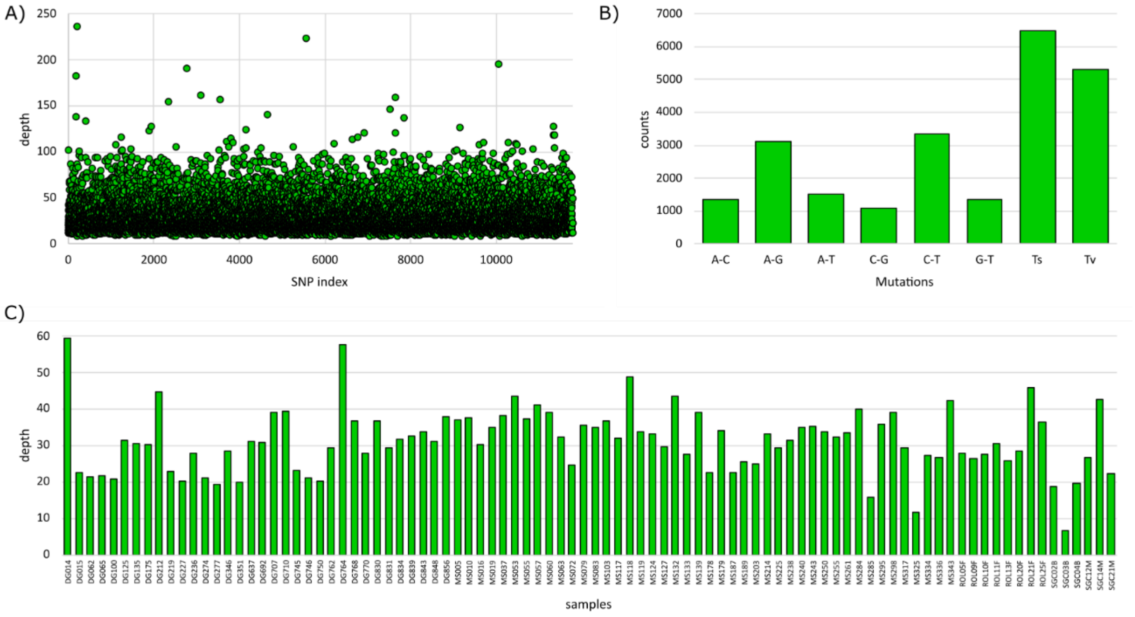

**Figure S2** – Quality metrics of the 11,782 SNP markers identified for 92 manioc (*Manihot esculenta*) varieties based on two double-digest genotyping-by-sequencing genomic libraries. A) mean sequence depth per locus, B) count of mutations (Ts = transitions, Tv = transversions), and C) mean sequence depth per sample.

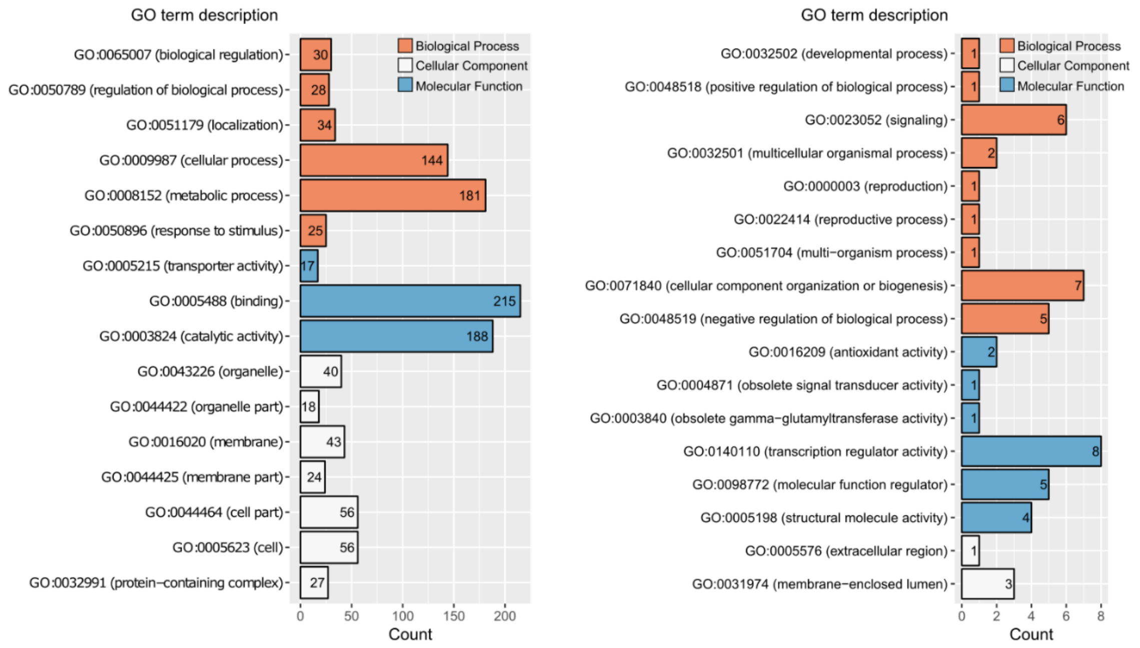

**Figure S3** – Summary of GO annotations of the 663 manioc predicted genes with SNPs putatively under selection. The annotations are grouped according to their biological processes, molecular functions, or cellular components. To facilitate visualization, bar plots were split for the GO terms occurring more than (left) or less than (right) 10 times. The number of genes with each annotation is indicated inside the bars.
